# The regenerative response of cardiac interstitial cells

**DOI:** 10.1101/2021.10.25.465720

**Authors:** Laura Rolland, Alenca Harrington, Adèle Faucherre, Girisaran Gangatharan, Laurent Gamba, Dany Severac, Marine Pratlong, Thomas Moore-Morris, Chris Jopling

## Abstract

Understanding how certain animals are capable of regenerating their hearts will provide much needed insights into how this process can be induced in humans in order to reverse the damage caused by myocardial infarction. Currently, it is becoming increasingly evident that cardiac interstitial cells play crucial roles during cardiac regeneration. To understand how interstitial cells behave during cardiac regeneration, we performed single-cell RNA sequencing (scRNA-seq) of regenerating zebrafish hearts. Using a combination of immunohistochemistry, chemical inhibition and novel transgenic animals, we were able to investigate the role of cell type specific responses during cardiac regeneration. This approach allowed us to identify a number of important regenerative mechanisms within the interstitial cell populations. Here, we provide here a detailed insight into how interstitial cells behave during cardiac regeneration and identify a number of novel features of these cells which will serve to increase our understanding of how this process could eventually be induced in humans.

## Introduction

The very limited regenerative potential of the adult mammalian heart underlies an increasing prevalence of heart failure(*1*). Studies using animal models such as the zebrafish and neonatal mice have shown that, following a substantial loss of myocardium, regeneration can be achieved through cardiomyocyte proliferation(*2, 3*). Furthermore, it has become increasingly evident that this process cannot occur without a suitable environment that is provided by multiple interstitial cell populations. Notably, following injury, clearing of cellular debris, neovascularisation and extracellular matrix scaffold constitution require the highly coordinated activity of interstitial cells such as immune cells, endothelial cells and fibroblasts(*4, 5*).

Recent studies have harnessed the power of single-cell level analysis to overcome difficulties associated with heterogeneous cell populations such as cardiac fibroblasts and macrophages. This has provided a more detailed overview of interstitial cell function after cardiac injury in adult mice and during cardiac regeneration in neonates(*6, 7*). Although the regenerating neonatal mouse heart is highly relevant for identifying mechanisms that could help promote adult human heart regeneration, it is also actively remodeling when subjected to injury, meaning certain features relevant to achieving regeneration in the adult heart may be missing. The zebrafish represents a complementary model for exploring cardiac regeneration as quiescent adult zebrafish myocardium is able to regenerate following significant injury(*2, 8*).

To further understand the process of cardiac regeneration in adult zebrafish, we have performed single cell analysis of interstitial cell populations in regenerating hearts. Furthermore, we provide a rigorous quantification of the different cell types present in the zebrafish ventricle, including cardiomyocytes, endothelium, epicardium, fibroblasts, macrophages and erythrocytes. Our analysis has revealed intriguing properties of fibroblasts, endothelial cells and macrophages that support cardiac regeneration in adult zebrafish.

Fibroblasts are involved in multiple processes associated with the cardiac response to injury and have previously been shown to play a crucial role during cardiac regeneration in zebrafish(*9-11*). Our data indicates many similarities in the injury response between zebrafish cardiac fibroblasts and their adult mammalian counterparts, however, we have also identified significant differences, most notably a disparity in myofibroblast gene expression. Endothelial cells make up the bulk of the cardiac interstitial population. Recent studies have determined that endothelial neovascularisation of the wound area is key for cardiac regeneration. This process lays down a framework over which the regenerating myocardium can be formed(*5*). Here we have determined that *tal1*, a gene essential for endocardial development, is required for cardiac regeneration in adult zebrafish. The macrophage response during cardiac regeneration also plays a pivotal role in ensuring a successful outcome(*12-15*). For a long time, the balance between inflammation and regeneration has been regarded as one of the key elements of this process. Interestingly, our data indicates that the resident macrophages present in the uninjured adult zebrafish heart appear to display an inflammatory M1 signature, which, following injury, is rapidly attenuated in conjunction with the arrival of recruited M2 macrophages. Furthermore, we have also determined that the matrix metalloproteinase, *mmp14*, is primarily expressed by recruited M2-like macrophages and plays a crucial role in allowing them to invade the damaged tissue.

Our study underlines the importance and variety of interstitial cell functions that support adult zebrafish heart regeneration. Furthermore, we have also compared our findings with published single cell RNA sequencing (scRNA-seq) studies of the interstitial cellular response in regenerating neonatal mice and non-regenerating adult mice following myocardial infarction(*6, 7*). In so doing we have endeavored to provide a balanced overview of the similarities and differences between regenerating and non-regenerating models.

## Results

### Single cell sequencing of regenerating zebrafish ventricle

In order to analyse the regenerative response of different interstitial cell populations, we adopted a scRNA-seq strategy. Following optimization of dissociation conditions and FACS-sorting of viable nucleated cells, we performed scRNA-seq (10x Chromium) of uninjured, sham operated and amputated (3 days, 7 days and 14 days post-amputation (dpa)) adult zebrafish ventricles. Altogether, after quality control, we obtained 18,739 transcriptional profiles and found that samples from uninjured and sham-operated zebrafish were largely comparable (Fig.1.A and Suppl Fig.1.A, Suppl Table I). Unbiased clustering of cells from uninjured and amputated hearts revealed 14 clusters comprising 9 distinct cell types (Fig.1.B, Suppl Fig 1.B and Suppl data.1 and 2). Major cell types included *cdh5*^+^ endothelium, *mpeg1.1^+^* myeloid cells (macrophages), *tcf21^+^* epicardium/epicardium-derived cells (EPDCs), *hbaa1^+^* erythrocytes and *sla2^+^* lymphoid cells (Fig.1.B, C). The latter included *lck^+^* T-lymphocytes and *pax5^+^* B-lymphocytes (Suppl Fig.1.C). Smaller populations included *mpx^+^* neutrophils, *itga2b^+^* thrombocytes and *sox10^+^* cardiac neural crest (and *rspo1^+^* neural crest derivatives) (Fig.1.B, C). Although we could detect a small population of *myl7^+^* cardiomyocytes, these cells were largely absent from our data set presumably because their size was not compatible with our scRNA-seq pipeline (Fig.1.B, C). Very rare “hybrids” i.e. expressing markers of multiple lineages, formed protrusions from cell-type specific clusters aimed in the direction of another cell type, as most obviously observed with the myocyte cluster (Fig.1.B). Cell types were clearly separated in the UMAP plots, with an exception of the myeloid and lymphoid lineages. These populations included converging sub-populations characterized by a high expression of cell-cycle related genes (e.g. *cdk1, mik67)* (Fig.1.C-E and Suppl data.1 and 2).

**Figure 1.**
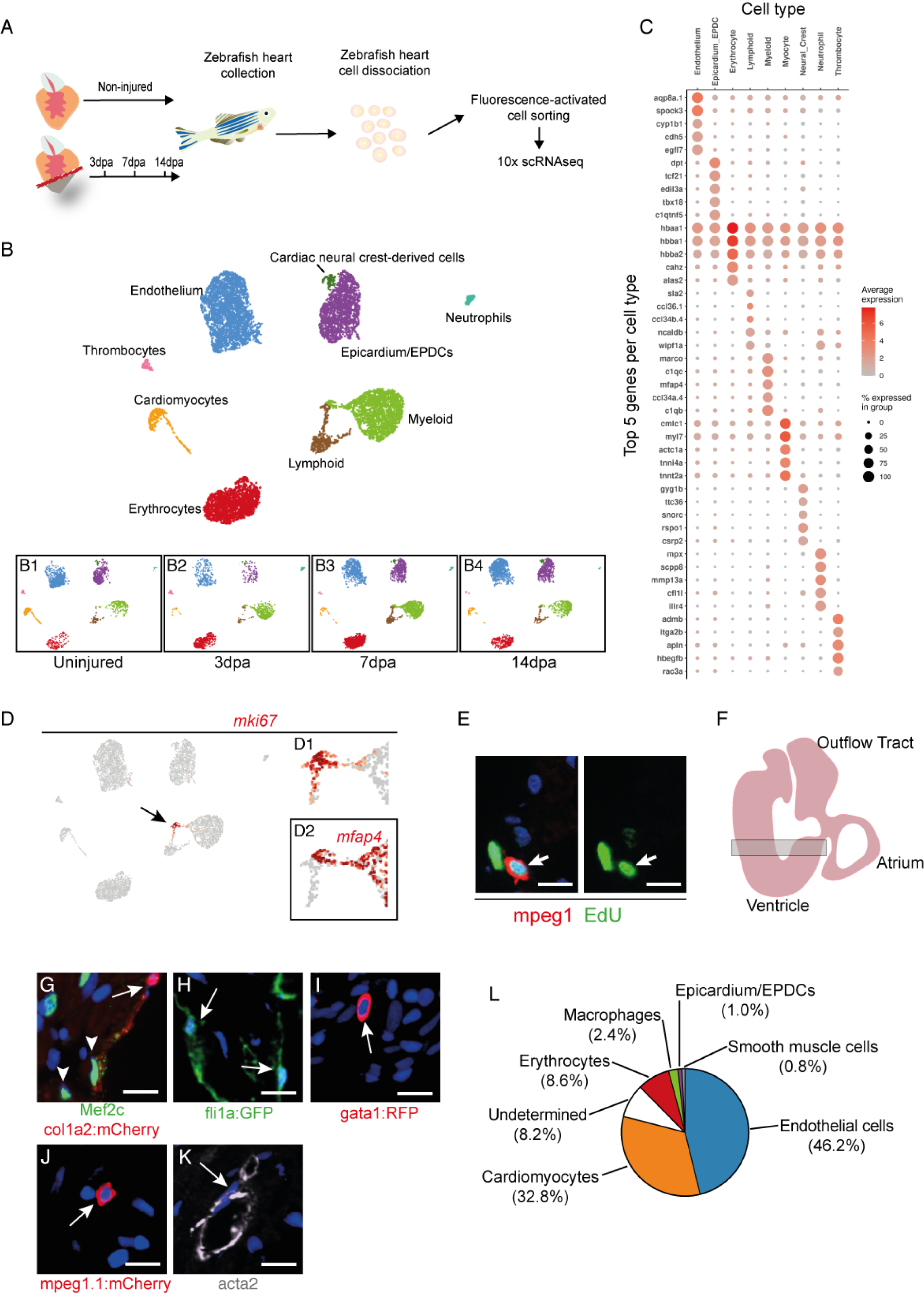
Single cell RNAseq analysis of regenerating zebrafish hearts. (**A**) Schematic of our scRNA-seq pipeline. (**B**) UMAP clusters of the different populations of cells identified in zebrafish hearts. (**B1-4**) UMAP clusters in uninjured, 3 days post amputation (dpa), 7 dpa and 14 dpa. (**C**) A DotPlot showing the 5 genes used to characterize the different cell types. (**D**) UMAP plot indicating the cluster of proliferating cells based on the expression of *mki67*, black arrow points to the cluster which is shown at higher magnification (**D1**). (**D2**) The proliferating cluster contains *mfap4^+^* myeloid (macrophages) and *mfap4^-^* lymphoid cells. (**E**) IHC of EdU labelled 3dpa regenerating zebrafish hearts. Mpeg1 (red) labels macrophages, EdU (green) labels proliferating cells, DAPI (blue) (scale bar 10μm). (**F**) Diagram of an adult zebrafish heart, the shaded rectangle indicates the area which was used to count the different cell type. (**G-K**) Examples of IHC images used to count the different cell types. Cardiomyocytes, Mef2c (green). Fibroblasts, Col1a2:mCherry (red). Endothelium, Fli1a:GFP (green). Erythrocytes, Gata1:RFP (red). Macrophages, Mpeg1.1:mCherry, (red). Smooth muscle cells, Acta2 (red). (scale bar 10μm). (**L**) Pie chart representing the proportions of different cell types present within the uninjured adult zebrafish ventricle. The average percentage of cells positive for each cell-type specific marker were identified among at least 800 cells per heart, with 5 hearts analysed for each marker.

Loss of large and/or fragile cells, most notably cardiomyocytes and endothelial cells, can lead to underrepresentation of these cell types in scRNA-seq datasets(*6*). Accurate evaluation of the relative proportions of these cell types, in particular at baseline, is essential for contextualizing cell-type specific responses to injury. Previous studies have established that the adult zebrafish heart is mainly composed of cardiomyocytes and endothelium(*16*), but the proportions of other key cell types, including resident macrophages and epicardium/EPDCs has not previously been described. Using IHC and several reporter lines, we were able to assign a cell-type identity to 91.8% of DAPI^+^ nuclei in the ventricle (Fig.1.F-L). These included Mef2c^+^ cardiomyocytes (32.8%), *Tg(fli1a:GFP)y1^+^* endothelial cells (46.2%), *Tg(gata1:dsred)^+^* erythrocytes (8.6%), *Tg(mpeg1.1:mCherry)^+^* macrophages (2.4%), *Tg(col1a2:mCherry)*^+^ epicardium/EPDCs (1%) and a-smooth muscle actin (Acta2)^+^ smooth muscle cells (0.8%) (Fig.1.G-L). It was essential to clearly identify and quantify zebrafish erythrocytes as they are nucleated in this species. We were unable to assign an identity to 8.2% of nuclei that may include cells with a weak reporter/IHC signal and rare populations such as lymphocytes or CNC-derived cells. Hence, as in the adult mouse heart, the endothelial and cardiomyocyte populations represented the most abundant cell types(*17*). However, the uninjured adult zebrafish heart presented a relatively low number of epicardium and epicardial-derived cells (EPDCs) such as fibroblasts. Indeed, we evaluated that epicardium/EPDCs represented 1% of the cells in adult the zebrafish heart. In comparison 15% of the adult mouse heart is comprised of fibroblasts(*17*).

### Macrophages

Our analysis revealed 3 macrophage clusters (MC1-3), including MC1 that was predominant in uninjured hearts. The top gene associated with these resident MC1 macrophages *versus* other myeloid clusters was *cxcr3.3,* a ligand scavenging receptor associated with reduced macrophage mobility(*18*). These cells also expressed the highest levels of markers of activated M1 macrophages (*tnfa, il1b, cd40, il6r*) and neutrophil-recruiting chemokines (*cxcl8a, csf3b*) (Suppl Fig.1.D and Suppl data.1). MC2 was predominant at 3dpa, and expressed high levels of genes associated with M2-like properties, including *ctsc* and *c1qa* (Suppl Fig.1.D and Suppl data.1). Also, MC2 macrophages expressed genes associated with recruited macrophages such as *ccr2*(*19*) and *apoeb*(*20*) as well as tissue healing including the copper chaperone *atox1*(*21*) (Suppl Fig.1.D and suppl data.1). MC3 was characterized by a very strong cell-cycle related gene signature, including *mki37, top2a, pcna* and *cdk1* (Fig.1.D, D1, D2 and Suppl data.1). In support of this, we were able to directly observe proliferating EdU-labelled macrophages in regenerating ventricles by immunohistochemistry (IHC) analysis (Fig.1.E).

### Endothelium

Based on cell counts, we determined that 46.2% of cells in the ventricle were endothelial/endocardial. Interestingly, our scRNA-seq data clearly showed that, both at baseline and following resection, endocardial endothelium had both an endothelial (*cdh5*) and mesenchymal (*pdgfra, col1a2*) signature, as previously observed in mouse(*22*). We obtained four endothelial cells clusters, all showing strong expression of pan-endothelial markers *aqp8a.1, cdh5* and *vwf* (Suppl Fig.1.B and Suppl data.2). Endothelial clusters (EC) 1 and 2 were abundant in control hearts. EC1 was characterized by expression of relatively high levels of collagen (*col1a1a/b, col1a2)* (Suppl Fig.1.E and Suppl data.1). EC2 was characterized notably by *nppc*, a key regulator of vascular homeostasis(*23*) whose receptor *npr3* was expressed by EPDCs at baseline (Suppl Fig.1.E and Suppl data.1). EC3 and EC4 increased in size over time following amputation (Fig.1.B1-B4). EC3 expressed relatively high levels of several heme-binding genes (*hbba1, hbba2, hbaa1, hbaa2*), albeit at a far lower level than erythrocytes (Fig.1.C and Suppl Fig.1.E and Suppl data.1 and 2.). Markers of venous endothelium, such as *kdrl*, were not specific to any EC cluster (Suppl Fig.1.E). Markers of lymphatic endothelium *prox1a* and *lyve1b* were expressed by a small subset of endothelial cells that did not segregate to any specific cluster (Suppl Fig.1.E and Suppl data.1).

### Epicardium/EPDCs

Within our dataset we could clearly delineate two Epicardial/EPDC clusters. Epicardium/EPDC cluster 1 (FB1), was characterized by a high expression of genes associated with extracellular matrix organization (*adamstl2, col18a1b)* and integrin binding (*edil3a, hapln1a*) (Suppl Fig.1.F and Suppl data.1). Cells in FB2 were notably characterized by a high expression of genes involved in complement activation (*c4, c4b, c6*) (Suppl Fig.1.F and Suppl data.1). Proportionately, FB1 was most abundant in unamputated hearts, whereas cell numbers in FB2 increased following injury (Fig.1.B1-B4). We were able to clearly identify epicardium and epicardial-derived fibroblasts based on their expression of the previously described fibroblast specific gene *tcf21*(*24*)(Fig.1.C and Suppl data.1 and 2.). Clustering did not seem to reflect epicardial *versus* epicardium derived cells. For example, *aldh1a2^+^* and *clu^+^* epicardial cells were present in subsets of both FB1 and FB2 (Suppl Fig.1.F and Suppl data.1).

### Neural crest

We were also able to identify a *sox10^+^* population of cardiac neural crest (CNC)-derived cells which were assigned to the FB2 cluster, probably because of relatively low numbers of cells (Suppl Fig.1.B). We classified these cells as CNC derived based on the high expression of markers of neural crest cells (*sox10^+^*, *pax3a^+^*, *apoda.1^+^*) and CNC-derived mesenchyme (*rspo1^+^*, *gyg1b^+^, csrp2^+^, hand2^high^*) (Fig.2.A and Suppl data.1 and 2.). Strikingly, these cells did not express the epicardium/EPDC-specific genes, *tcf21* and *tbx18* (Fig.2.A) nor the FB2-specific gene, *adh8a* (Suppl data.1). Within the CNC cluster, we could clearly distinguish *sox10^high^;rspo1^-^* and *sox10^low^*;*rspo1^+^* sub-clusters. Previous studies in zebrafish have shown that *sox10*-expressing cells are concentrated in the atrioventricular valves(*25*). In mice, *Rspo1* has recently been reported to be expressed in epicardial cells(*7*). IHC analysis of neonatal mouse hearts confirmed that RSPO1 was indeed associated with the epicardial layer (Suppl. Fig 2.A). Similarly, IHC revealed that *rspo1*^+^ cells were also most abundant in the zebrafish ventricle epicardial layer (Fig.2.B). However, re-analysis of the scRNA-seq data by Wang *et al*.(*7*) confirmed that, in contrast to what we observed in zebrafish, murine *Rspo1^+^* cells were clearly expressing epicardial markers such as *Tcf21* and *Tbx18* (Suppl Fig 2.B), underlining a divergence in the profiles of *Rspo1^+^* cells in these species.

**Figure 2.**
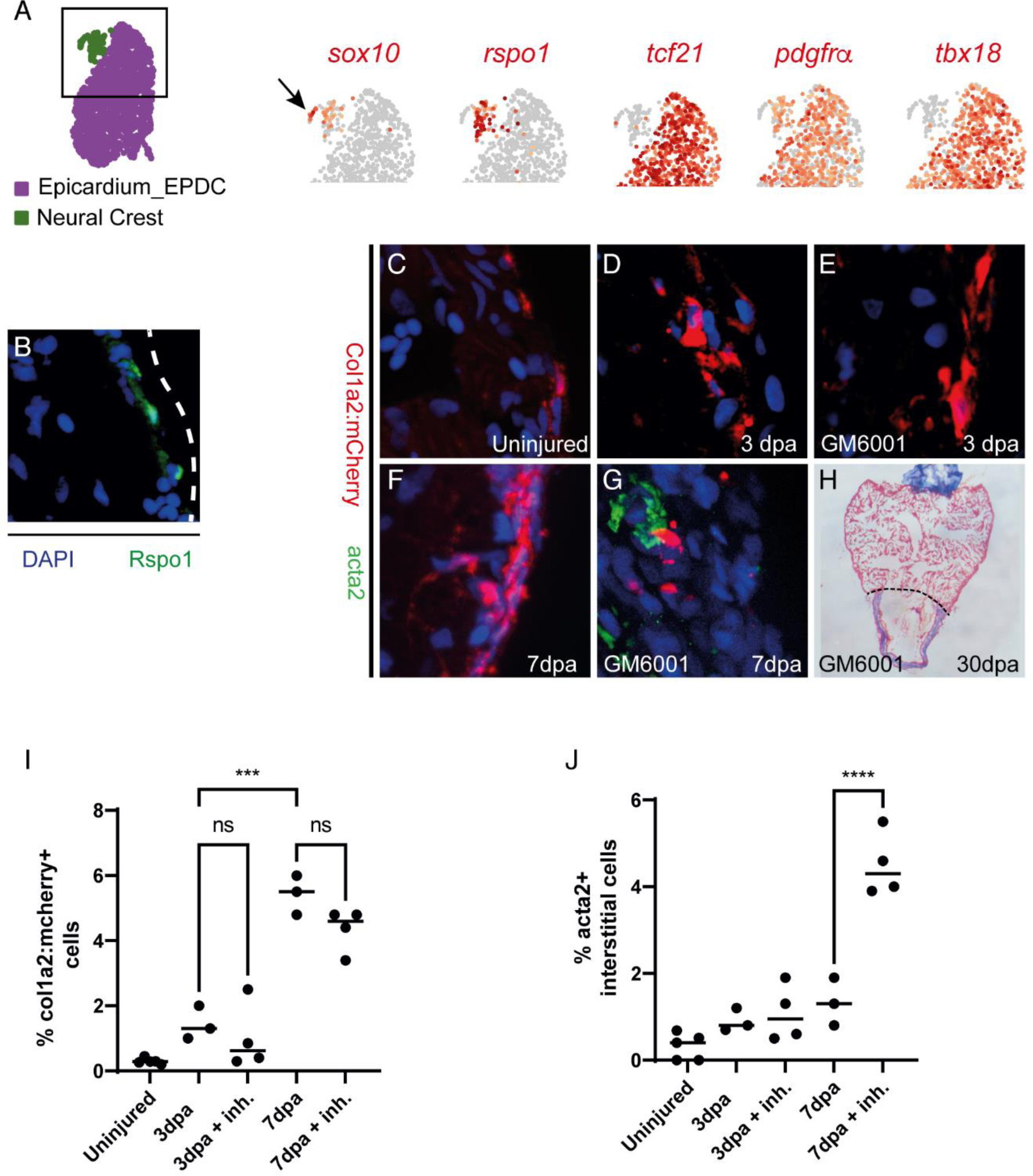
Mesenchymal lineages and cardiac fibroblast activation in the regenerating zebrafish heart. (**A**) UMAP plot indicating the epicardial (purple) and neural crest (green) clusters. The black square delineates the area depicted in the subsequent UMAP plots. UMAP plots depicting the relative expression of *sox10* (Neural crest) (black arrow indicates the neural crest cluster), *rspo1*, *tcf21* (Epicardium/EPDC), *pdgfra* (Epicardium/EPDC) and *tbx18* (Epicardium/EPDC). Note that *rspo1* is restricted to the neural crest cluster and does not segregate with either *tcf21* or *tbx18*. (**B**) IHC image of a *Tg(col1a2:mCherry)* zebrafish heart labelled with a RSPO1 antibody (green). (**C-G**) IHC images of *Tg(col1a2:mCherry)* zebrafish hearts showing Col1a1:mCherry (red, fibroblasts) and Acta2 (green) in uninjured (**C**), 3dpa (D), 3dpa inhibitor treated (**E**), 7dpa (F) and 7dpa inhibitor treated hearts (**G**). Note the absence of Col1a2:mCherry and Acta2 co-localisation. Although Acta2 (green) is present in the 7dpa inhibitor treated sample this does not co-localise with Col1a2:mCherry (**H**) AFOG staining of a MMP inhibitor (GM6001) treated zebrafish heart at 30dpa, black dashed line indicates the plane of amputation. Note the presence of a large fibrin (red)/collagen(blue) scar. (**I**) Graph showing the percentage of cells, from an average of 650 cells per heart, that were Col1a2:mcherry^+^ in the different experimental conditions. (**J**) Graph showing the percentage of interstitial cells, from an average of 650 cells per heart, that produced Acta2 in the different experimental conditions. Note that inhibitor treatment significantly increases the number of Acta2 cells. P values were calculated by 1-way ANOVA. \*\*\**P<0.001*, **** *P<0.0001*; ns, non-significant.

### A hallmark of activated mammalian cardiac fibroblast is absent in zebrafish

Following injury, we observed a strong fibrotic response, which was initiated in epicardium/EPDCs, notably with a robust upregulation of *periostin* (*postnb)* and *fibronectin* (*fn1a/b*) expression(*26*) (Suppl Fig.2.C and Suppl data.1 and 2). In contrast, *acta2,* a gene associated with injury-induced myofibroblast activation in mammals(*26*), was not upregulated at any time point in the epicardium/EPDC lineage (Suppl Fig.2.C). To confirm this observation we performed cardiac amputations on *Tg(col1a2:mCherry)* zebrafish followed by IHC for Acta2 at 3dpa and 7dpa (Fig.2.C-J). Surprisingly, although the number of Col1A2:mCherry^+^ cells increased following injury, this was not accompanied by an increase in Acta2 expressing cells (Fig.2.D,I,J). Elevated Acta2 production can be observed in non-regenerating adult mammalian hearts following injury and, similarly, an increase in Acta2 production has also been observed in adult zebrafish mutants which are unable to regenerate their hearts(*27*). Based on these observations we surmised that inhibiting cardiac regeneration in adult zebrafish may also lead to an increase in Acta2 production and allow us to further investigate whether or not this was associated with fibroblasts. The pan– MMP inhibitor GM6001 has previously been reported to significantly inhibit zebrafish cardiac regeneration(*28*). In agreement with this we also found that treating adult zebrafish with GM6001 following apical resection resulted in a failure to regenerate at 30dpa and the formation of a large fibrin/collagen scar (Fig.2.H). Using this protocol we performed cardiac amputations on *Tg(col1a2:mCherry)* zebrafish followed by IHC for Acta2 in the lower ventricle at 3dpa and 7dpa (Fig.2.E,G,I,J). Interestingly, inhibiting cardiac regeneration with GM6001 led to a significant increase in the number of interstitial Acta2^+^ cells at 7dpa (Fig.2.G,J). However, Acta2 did not co-localise with *col1a2:mCherry*^+^, indicating that cell types other than fibroblasts were upregulating Acta2 expression (Fig.2.E,G). This is in agreement with a recent study that reported the expression of smooth muscle-specific genes outside of the epicardial lineage, notably in the endocardium(*29*). To determine whether the expression of *ACTA2* by fibroblasts varies between species we re-analysed 2 previously published scRNA-seq datasets from non-regenerating adult mice and regenerating neonatal mice after myocardial infarction(*6, 7*)(Suppl Fig.2.C). Interestingly, the sustained expression of the fibrosis associated gene, *Postn,* 3 days after injury appeared remarkably similar between adult mice and zebrafish (Suppl Fig.2.C). On the other hand, following myocardial infarction in adult mice and P8 neonates there was a robust upregulation of *Acta2* expression by fibroblasts 3 days post injury, a feature which was absent in the adult zebrafish after cardiac injury (Suppl Fig.2.C). The picture in regenerating P1 neonate fibroblasts was far less clear as they already expressed *Acta2* at baseline along with high levels of *Postn,* presumably because the P1 neonatal heart is still undergoing widespread remodeling (Suppl Fig.2.C). Taken together, our data indicates that collagen-producing cardiac fibroblasts in adult zebrafish do not upregulate Acta2 in response to injury.

### Tal1 is a regulator of the endothelial regenerative response

The zebrafish endothelial response to injury involves a rapid change in gene expression followed by wound neovascularization, a process which lays the foundation for the proliferating cardiomyocytes to regenerate the missing myocardium(*5*). A number of genes have been directly linked to the endothelial regenerative response in zebrafish such as *vegfaa*, *aldh1a2*, and *notch1b*(*30-32*). We could not detect any significant upregulation of any of these genes, which could be due to rapid changes in their expression occurring outside of the time points analysed here, as reported for *vegfaa*(*30*). On the other hand, although the average level of expression of *notch1b* in the endothelium did not change significantly we could detect an increase (8%) in the proportion of endothelial cells expressing this gene at 3dpa compared with uninjured controls. The re-expression of developmental genes is also a hallmark of regenerating endothelium(*5*). We found that the proportion of cells expressing genes required for endothelial development increased during cardiac regeneration (*foxc1a, foxc1b,* (Suppl Fig 3.A,B)(*33*)). The BHLH transcription factor Tal1 is also essential for endocardial development and for maintaining endocardial identity, in particular Tal1 is required for establishing endothelial Tjp1 tight junctions(*33*). Previous research has shown that *tal1* is upregulated during cardiac regeneration in zebrafish(*34*). In agreement with this, our scRNA-seq analysis indicates that there was an increase in the proportion of *tal1* expressing endothelial cells during regeneration (Fig.3.A,B). This increase peaked at 7dpa before returning to pre-injury levels by 14dpa (Fig.3.B). Without detailed lineage tracing we are presently unable to determine whether the increase in *tal1* expressing endothelial cells was due to the proliferation of existing *tal1* expressing cells or because more endothelial cells have begun producing *tal1* de novo. To confirm our scRNA-seq data, we performed IHC for Tal1 on adult *Tg(fli1a:GFP)y1* zebrafish cardiac sections and were able to clearly observe Tal1 positive endothelial cells (Fig.3.C-E). Because Tal1 is an obligate dimer and can form complexes with a variety of proteins which will subsequently dictate which transcriptional programs to activate/deactivate(*35*), we extended our analysis to known co-factors of Tal1. LIM only 2 (LMO2), forms a multi-protein complex with TAL1 and directs it towards specific targets genes. Previous research indicates that in the absence of LMO2, TAL1 is able to target other genes for expression/repression(*35*). We found that *lmo2* was downregulated during the early stages of regeneration (Suppl Fig 3.C). In order to analyse this in more detail we reclustered *tal1^+^* endothelial cells to identify changes in gene expression associated with this sub-population (Fig.3.F). Interestingly, this analysis revealed that although *lmo2* expression was evenly distributed in 3 of the *tal1* positive (*tal1^+^*) clusters (T1,T2,T3) it was reduced in the fourth (T4)(Fig.3.G). Furthermore, the expression of the *Lmo2*^-^/*Tal1^+^* target gene *cgnl1*(*35*), a component of tight junctions and implicated in neovascularization, shows the highest level of expression in cluster T4 (Fig.3.H). We could also observe a downregulation of *lmo2* at 3dpa and 7dpa specifically in *tal1^+^* endothelial cells (Fig.3.I). Conversely, *cgnl1* became upregulated in *tal1^+^* cells at 7dpa (Fig.3.J). Although these observations were below the threshold of significance (P<0.05), this evident trend prompted us to target *tal1* directly to determine whether this core endocardial developmental gene could be involved in cardiac regeneration. In order to achieve this we generated a transgenic zebrafish line which can express a dominant negative Tal1 isoform specifically in endothelial cells following treatment with tamoxifen (*fliEP:Ert2CreErt2; fliEP:loxRFPlox:DNtal*) (Suppl Fig 4.A-G). The dominant negative Tal1 isoform lacks the basic DNA binding domain but is still able to form multi-protein transcription complexes and thus inhibit native Tal1 associated transcription(*36*). We first assessed whether this transgenic line was functional by inducing DN *tal* expression during early zebrafish development. This procedure caused cardiac developmental defects reminiscent of those previously described for *tal1* knockout zebrafish (Suppl Fig 4. E-G). To determine whether *tal1* is required for heart regeneration we induced expression of DN *tal* prior to cardiac resection. Histological staining of heart sections at 30dpa revealed that expression of DN *tal* inhibited the regenerative process, as evidenced by a large fibrin/collagen scar (n=5) (Fig.3.K-P.). Lastly, to determine whether the expression of *tal1* in the endothelium is zebrafish specific we re-analysed 2 previously published scRNA-seq datasets from non-regenerating adult mice and regenerating neonatal mice after myocardial infarction and observed that *Tal1* is present within the endothelial populations(*6, 7*) (Suppl Fig.4.H). Taken together these data indicate that *tal1* is a key regulator of the endothelial response during cardiac regeneration.

**Figure 3.**
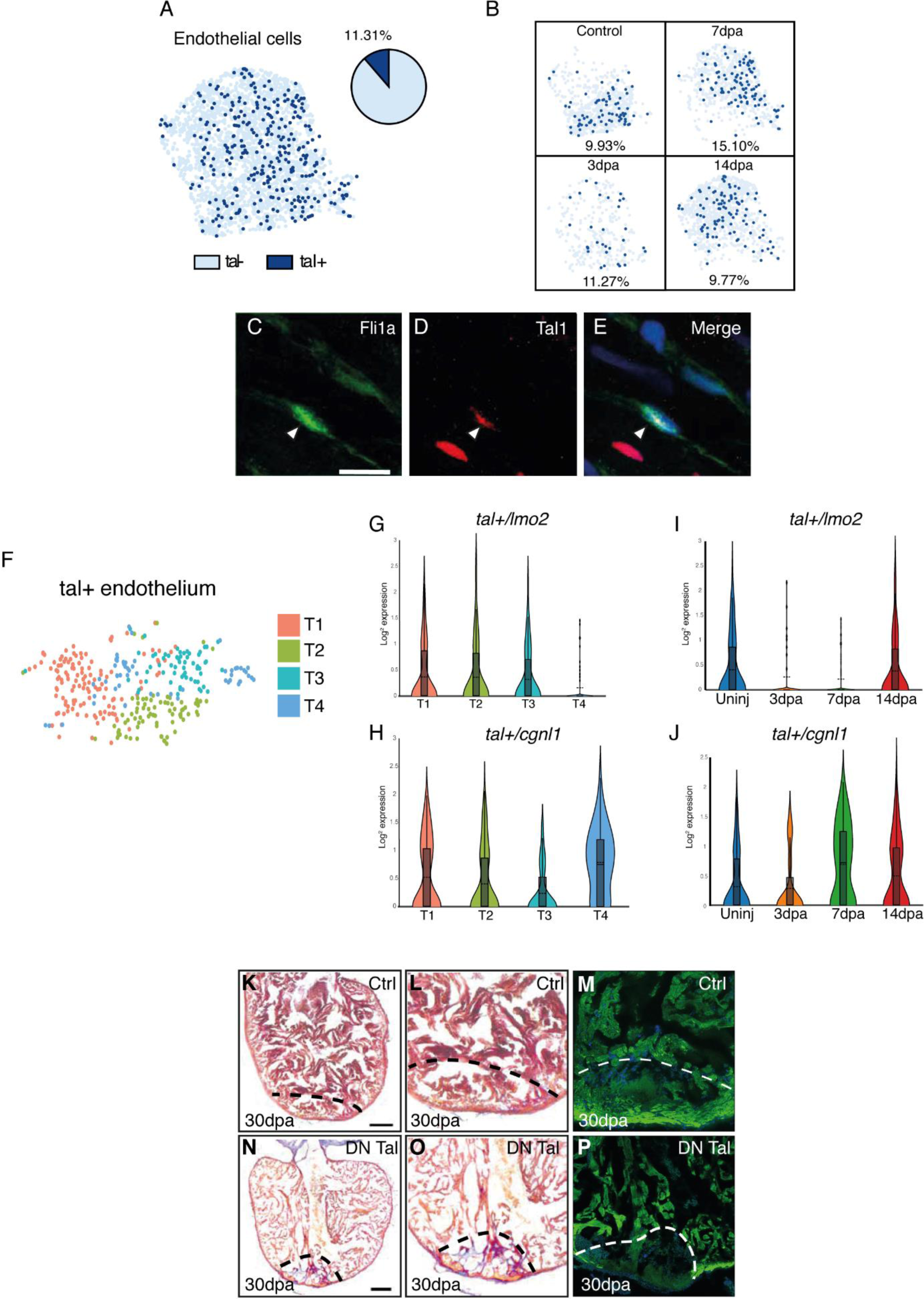
Tal1 is a regulator of the endothelial regenerative response. (**A**) UMAP plot of the endothelial cell cluster (light blue dots) showing *tal1* is expressed in a subset of these cells (dark blue dots). The pie chart indicates the total proportion of endothelial cells which express *tal1*. (**B**) UMAP plots of the endothelial population at different stages of regeneration (control/unamputated, 3dpa, 7dpa and 14dpa). The proportion of *tal1* expressing endothelial cells (dark blue dots) is given as a percentage underneath each UMAP plot. (**C-E**) IHC for endothelial cells (**C**, green), Tal1 (**D**, red), merged image (**E**) (scale bar 10μm). (**F**) Analysis of *tal1* expressing cells. UMAP plot of re-clustered *tal1* expressing cells. Coloured boxes represent the different clusters within this population (T1-T4) (**G-J**). Violin plots indicate the expression of either *lmo2* or *cgnl1* in each cluster (**G,H**) and at each time point during regeneration (**I,J**). (**K**) AFOG staining of a control heart at 30dpa, black dashed line indicates the plane of amputation (scale bar 200μm).(**L**) The same image at higher magnification. (**M**) IHC of the same heart at 30dpa labelled with an anti-α sarcomeric actin antibody (green). (**N**) AFOG staining of a DN Tal1 expressing heart at 30dpa, black dashed line indicates the plane of amputation (scale bar 200μm). (**O**) The same image at higher magnification, note the presence of a large fibrin (red)/collagen(blue) scar. (**P**) IHC of the same heart at 30dpa labelled with an anti-α sarcomeric actin antibody (green), note the large area beneath the white dashed line which is unlabeled indicating an absence of cardiomyocytes.

**Figure 4.**
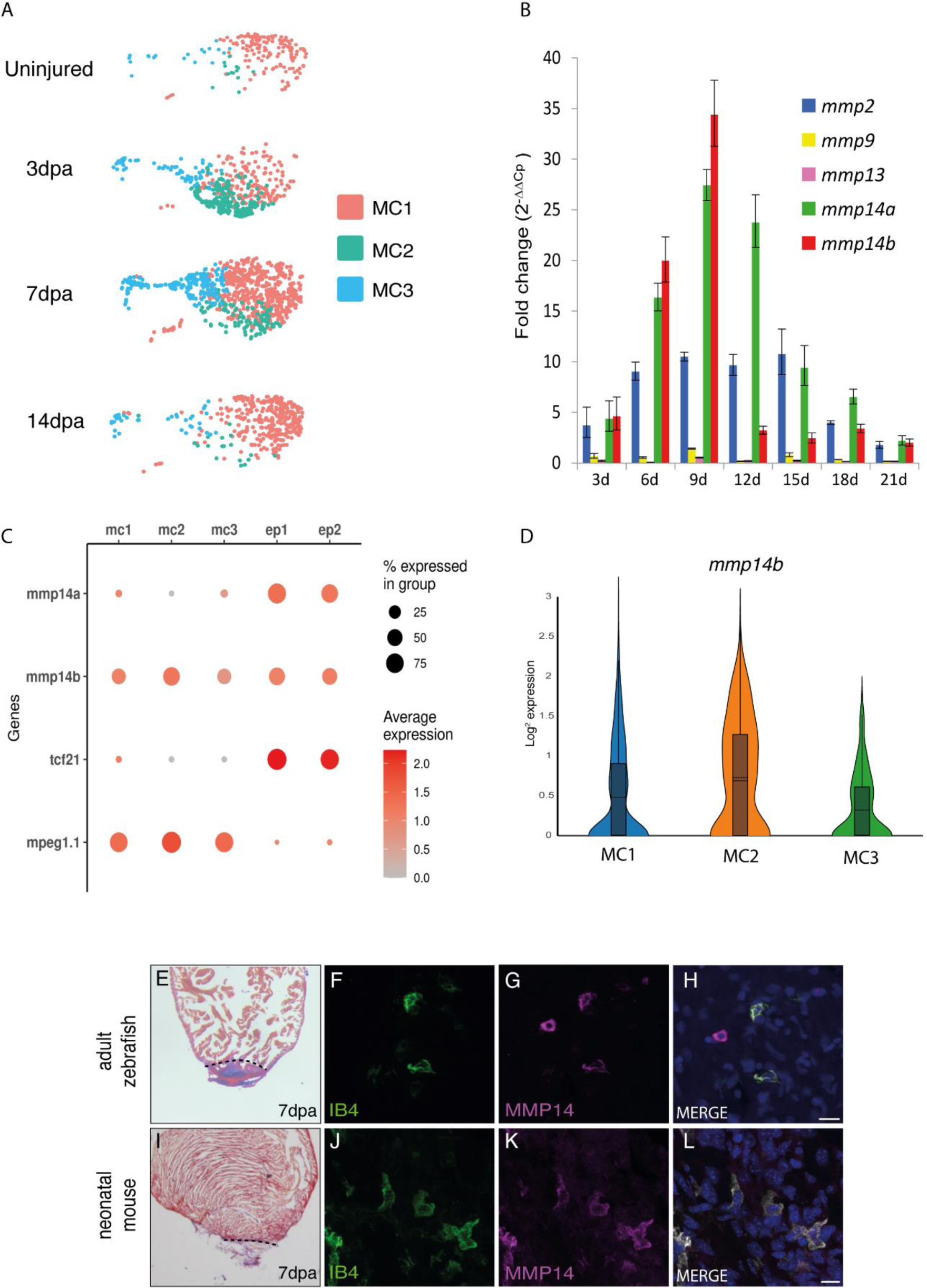
MMP14 is expressed by macrophages during cardiac regeneration. (**A**) UMAP plots of the macrophage population at different stages of regeneration (uninjured, 3dpa, 7dpa and 14 dpa). The coloured boxes indicate the different macrophage clusters (MC1-3). (**B**) RT-QPCR analysis of *mmp* expression during cardiac regeneration. (**C**) A DotPlot showing the relative expression levels of *mmp14a* and *mmp14b* in macrophages (MC1-3) and fibroblasts/epicardium (FB1,2). *tcf21* was used to identify the fibroblast population and *mpeg1.1* the macrophage population. (**D**) Violin plot comparing the expression of *mmp14b* in either the MC1, MC2 or MC3 macrophage clusters. (**E**) AFOG staining of a wildtype zebrafish heart at 7dpa, black dashed line indicates the plane of amputation. Note the presence of a large fibrin (red)/collagen(blue) scar. (**F-H**) IHC of a 7dpa zebrafish heart section for (**F**) IB4 (green, macrophages), (**G**) Mmp14 (magenta) and a merged image of F and G (scale bar 10μm)(**H**). (**I**) AFOG staining of a neonatal mouse heart section at 7dpa, black dashed line indicates the plane of amputation. Note the presence of a large fibrin (red)/collagen(blue) scar. (**J-L**). IHC on a neonatal mouse heart section at 7dpa (**J**) IB4 (green, macrophages), (**K**) MMP14 (magenta) and a merged image of J and K (**L**).

### MMP14 expressing macrophages are required for cardiac regeneration

Macrophages play a crucial role during cardiac regeneration(*15*). We were able to identify 3 clusters of macrophages within our scRNA-seq data set. As described earlier, MC1 represented a resident population of inflammatory/M1 macrophages, which was predominant in uninjured hearts (Fig.4.A). MC2 was comprised of recruited macrophages, which were abundant at 3dpa, persisted through 7dpa and resolved by 14dpa (Fig.4.A). This cluster was enriched for transcripts commonly associated with M2 polarised macrophages such as *cd9a/b*, inflammasome associated genes (*txnipa*, *caspa* and *atp13a2*) and *csf1ra*(*37-40*) (Fig.4.A and Suppl Fig.5 A-F and Suppl data.1). Furthermore, MC2 was also enriched for genes previously described to be associated with regenerating macrophages in adult zebrafish (*fabp11a*, *lgals9l1* and *lgmn*(*41*)) (Suppl Fig 5.G-I and Suppl data.1). Lastly, MC3 was enriched for genes associated with the cell cycle such as *pcna* and *mki67* indicating that macrophage proliferation also occurs during cardiac regeneration in zebrafish, similar to the proliferation observed in mouse hearts following myocardial infarction (MI)(*42*) (Suppl Fig 6.A,B and Suppl data.1).

**Figure 5.**
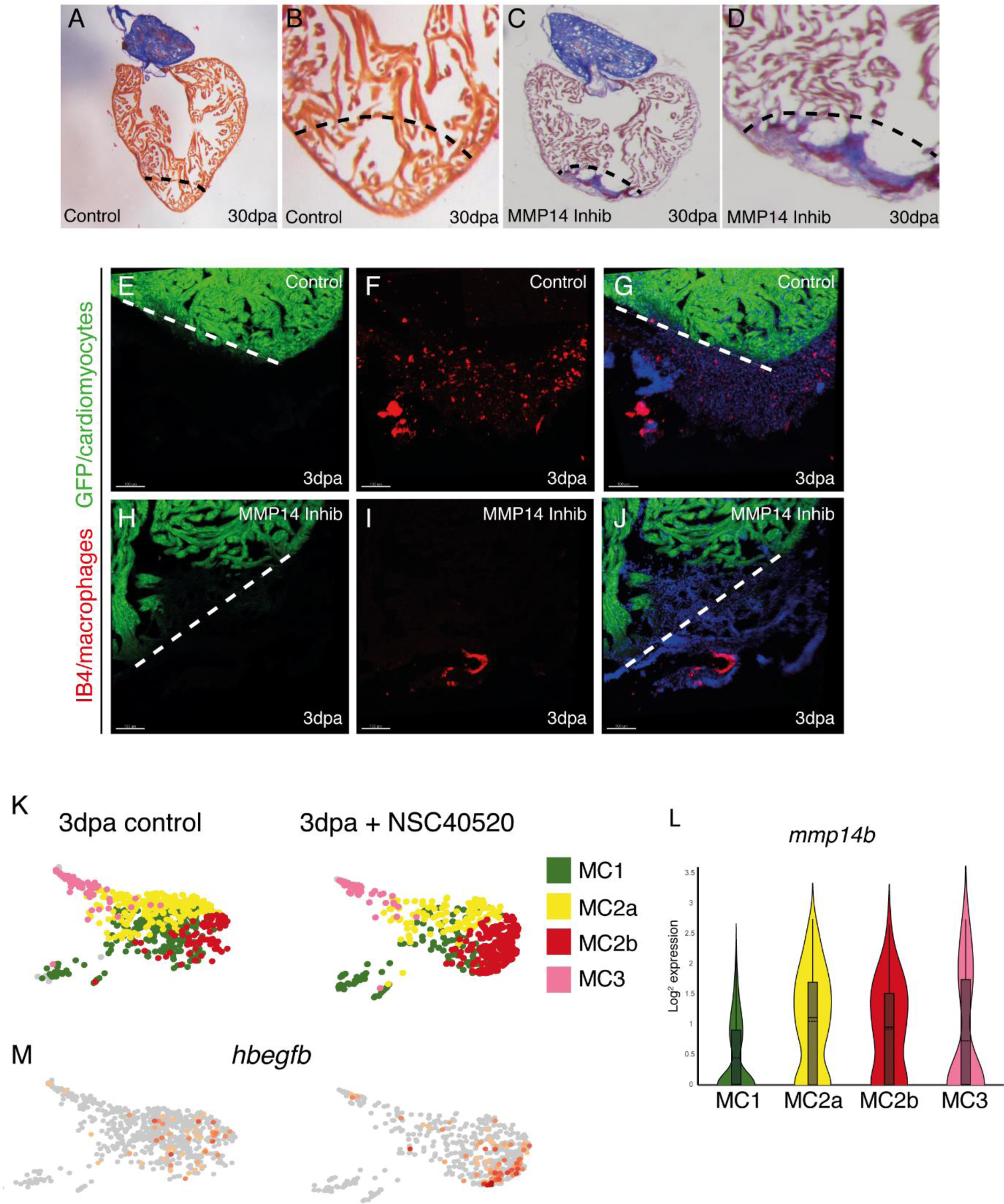
MMP14 expressing macrophages are required for cardiac regeneration. (**A,B**) AFOG staining of a control zebrafish heart at 30dpa, black dashed line indicates the plane of amputation (scale bar 200μm). (**C,D**) AFOG staining of a MMP14 inhibitor treated zebrafish heart at 30dpa, black dashed line indicates the plane of amputation (scale bar 200μm). Note the presence of a large fibrin (red)/collagen(blue) scar. (**E-G**) IHC on a 3dpa control zebrafish heart section for (**E**) cardiomyocytes (GFP, green) (scale bar 100μm), (**F**) macrophages (IB4, red) and a merged image of E and F (**G**). Note the presence of numerous macrophages in (**F**). (**H-J**) IHC on a 3dpa MMP14 inhibitor treated zebrafish heart section for (**H**) cardiomyocytes (GFP, green) (scale bar 100μm), (**I**) macrophages (IB4, red) and a merged image of H and I (**J**). Note the absence of macrophages in (**I**). (**K**) UMAP plots of the macrophage population in control and MMP14 inhibitor treated zebrafish hearts at 3dpa. The coloured boxes indicate the different macrophage clusters (MC1-3). (**L**) Violin plot comparing the expression of *mmp14b* in the MC1, MC2a, MC2b and MC3 macrophage clusters. (**M**) UMAP plots depicting the relative expression of *hbefgb* in control and MMP14 inhibitor treated macrophages. Note the increased expression of *hbefgb* in the MMP14 inhibitor treated recruited macrophage cluster (MC2b).

The expression profiles of *tnfa* and *il1b* are commonly used to distinguish between M1 and M2 macrophage polarization. We observed that the highest expression of *tnfa* was in uninjured resident macrophages (Suppl Fig 7.A). In agreement with this, we also observed that the expression of *txnipa*, a gene whose expression is negatively regulated by *tnfa*(*43*), was sharply upregulated at 3dpa as *tnfa* expression subsided (Suppl Fig 7.B). Furthermore, resident macrophages also expressed higher levels of *il1b* compared to the recruited MC2 population (Suppl Fig 1.D). Because this appears to be at odds with previous reports which indicate that recruited macrophages display a more pro-inflammatory signature than resident cells(*44, 45*), we re-analysed 2 previously published scRNA-seq datasets from non-regenerating adult mice and regenerating neonatal mice after myocardial infarction(*6, 7*)(Suppl Fig.8). Expression of the chemokine receptor Ccr2 has previously been shown to effectively identify recruited macrophages following myocardial injury(*19*). During cardiac regeneration in P1 neonatal mice there was an increase in macrophage *Ccr2* expression 1 day after injury, which is also associated with an increased expression of *Tnfa* and *il1b*, indicating a wave of pro-inflammatory macrophages are recruited to the heart shortly after injury (Suppl Fig.8). Similarly, in adult mice after MI there is an influx of *Ccr2* expressing macrophages at 3 days post injury which coincides with an increase in *Il1b* expression (Suppl Fig.8). Our own data indicates that during cardiac regeneration in adult zebrafish *ccr2* was expressed at low levels 3 days after injury, predominantly in the recruited MC2 population (Suppl Fig.8). Although it appears from our data that resident macrophages (MC1) expressed higher levels of *tnfa* and *il1b* than the recruited population (MC2) (Suppl Fig.8), we cannot rule out the possibility of a rapid influx of pro-inflammatory macrophages at an earlier timepoint, as observed in neonatal mice. In support of this notion, the expression profiles for *tnfa* and *il1b* at 3 days post injury followed a similar pattern in both neonatal mice and adult zebrafish. In particular there was a marked reduction in expression of both these genes at 3 days post injury compared to uninjured controls (Suppl Fig.8). Further analysis will be required at earlier timepoints to fully decipher the macrophage inflammatory response during cardiac regeneration in zebrafish.

Our data also reveals a potential zebrafish specific mechanism for modulating the response of macrophages to CXCL chemokine signals. In mammals, G protein coupled chemokine receptor 3 (*CXCR3)* signalling is required for recruiting macrophages to the site of injury/infection(*46*). Previous reports indicate that zebrafish possess 2 orthologs of this gene, *cxr3.2* and *cxcr3.3*. Although *cxcr3.2* appears to be a functionally active receptor, *cxcr3.3* lacks the ability to activate downstream signaling pathways and hence acts as a scavenger of *cxcr3* ligands, effectively dampening down the *cxcr3.2* response(*18*). Although all the macrophage clusters express *cxcr3.2*, *cxcr3.3* is particularly enriched in resident, MC1, macrophages which would presumably reduce their ability to respond to *cxcr3* ligands (Suppl Fig.9.A,B and Suppl data.1). Matrix metalloproteinases (MMP) are also well established players in wound healing and regeneration and have been linked to a variety of processes that are necessary to resolve damaged tissue^10^. Transcriptomic analysis of zebrafish heart regeneration has consistently identified an upregulated expression of various MMPs during this process(*28, 34, 47*). Furthermore, MMPs have been shown to be involved in zebrafish fin regeneration(*48*), newt limb regeneration(*49*) and salamander limb regeneration(*50*). We initially assessed MMP expression during cardiac regeneration by qPCR and observed a dynamic response during this process, in particular the expression of *mmp2* and *mmp14a/b* (zebrafish possess 2 *MMP14* orthologs) increased substantially (Fig.4.B). Analysis of our scRNA-seq data indicates that *mmp14a/b* were predominantly expressed by fibroblasts and macrophages. In particular *mmp14a* was restricted to the fibroblast population and was absent in macrophages while, *mmp14b* appears to be expressed by both populations (Fig.4.C and Suppl data.1). Of particular interest, we also observed that *mmp14b* is predominantly expressed by the recruited MC2 macrophages (Fig.4.D and Suppl data.1). To confirm that macrophages express MMP14 during regeneration we performed IHC for MMP14 on regenerating zebrafish and P1 neonatal mouse hearts (7dpa). In this manner, we were able to clearly detect MMP14 positive macrophages present during regeneration in both species (Fig.4.E-L). To gain further insight into MMP14 expression by macrophages we re-analysed 2 previously published scRNA-seq datasets from non-regenerating adult mice and regenerating neonatal mice after myocardial infarction(*6, 7*)(Suppl Fig.10.A-C). Interestingly, *Mmp14*^+^ macrophages were absent at baseline and three days post MI in both adult and neonatal mice (Suppl Fig.10. B, C). This is in contrast with zebrafish, in which *mmp14b* expressing macrophages were present in uninjured hearts and peak at 3dpa (Supp Fig.10.A). To determine whether Mmp14 is required for cardiac regeneration we employed a specific inhibitor of this protein, NSC40520, which blocks the collagenolytic activity of MMP14 but not its ability to activate other targets such as MMP2(*51*). Histological analysis of cardiac sections taken from 30dpa adult zebrafish indicated that NSC40520 treatment significantly impaired cardiac regeneration, resulting in the formation of a fibrin/collagen scar (n=5) (Fig.5.A-D). To determine whether Mmp14 inhibition disrupts the macrophage regenerative response, we repeated these experiments at 3dpa, a time point when M2/Mmp14^+^ recruited macrophages were most abundant. IHC analysis of cardiac sections revealed substantial numbers of macrophages in the wound region at 3dpa in untreated control samples (n=5) (Fig.5.E-G). Strikingly, macrophages were noticeably absent in the wound region of NSC40520 treated samples (n=5) (Fig.5.H-J).

Based on these observations, we performed scRNA-seq analysis to assess what effect Mmp14 inhibition had on the macrophage population. In order to identify the early events affected by NSC40520 treatment we focused on 3dpa when recruited *mmp14b* expressing macrophages are most abundant. Analysis of the fibroblast population within this dataset indicates that overall there are no dramatic changes in the expression of genes associated with the fibrotic response following inhibitor treatment (Suppl Fig.11.A-D). However, the expression of *mmp14a* and *mmp14b* did increase, which may reflect an attempt by these cells to counteract the Mmp14 inhibition (Suppl Fig.11.E, F). Within the macrophage population we could identify 2 clusters of recruited macrophages, MC2a and MC2b which were enriched for *mmp14b* and *cd9b* (Fig.5. K, L and Suppl Fig.12.A, B). Furthermore, we were able to identify a number of genes whose expression was significantly upregulated in *mmp14b* enriched macrophages following inhibitor treatment. Notably we observed a robust upregulation of *fabp11a* and *lgals9l* expression, both of which have previously been associated with regenerating macrophages(*41*)(Suppl Figs.12.C, D). Of potential interest the heparin-binding epidermal growth factor-like growth factor, *hbegfb,* which is cleaved and activated by MMP14(*52*), is specifically upregulated in recruited macrophages following inhibitor treatment (Fig.5.M and Suppl Fig 12.E). Taken together our data indicates that the macrophage response to cardiac injury in adult zebrafish is reminiscent of the early response observed in neonatal mice after MI. Furthermore, we have determined that Mmp14 is required for effective cardiac regeneration and that inhibiting the collagenolytic activity of Mmp14 results in defective migration of M2 recruited macrophages into the wound region.

## Discussion

Here we report a detailed scRNA-seq driven analysis of how interstitial cells behave during cardiac regeneration in zebrafish. Our data has uncovered a number of intriguing insights into the regenerative process and also highlights notable differences between regenerating and non-regenerating species. Regarding the cellular composition of the adult zebrafish heart, when compared to the adult mouse heart we found largely comparable proportions of endothelial cells and cardiomyocytes. However, we observed a notable difference in the proportion of cells derived from the epicardial lineage. In particular, the number of fibroblasts present in the adult zebrafish heart (1%) was considerably lower than what has been reported for the adult mouse heart (15%)(*16*). In mammals, loss of cardiac tissue due to myocardial infarction (MI) triggers the differentiation of fibroblasts into myofibroblasts which are involved in replacing the lost tissue with a fibrotic scar(*26*). Although this initial phase is essential for maintaining cardiac integrity, the fibrotic response can spread throughout the heart leading to impaired cardiac function and ultimately failure. Indeed, it has been proposed for some time that the difference between scarring and regeneration could be influenced by the fibrotic response to injury(*53*). One feature of this response involves *α Smooth Muscle Actin* (*ACTA2*), a gene commonly restricted to smooth muscle cells, becoming specifically upregulated in myofibroblasts(*26*). Our data, combined with re-analysis of scRNA-seq data from mouse models, showed that, following injury, the epicardial/fibroblast lineage strongly upregulated a number of myofibroblast-associated genes such as *Postn/postnb* and *Fn1*/*fn1a* in both species. However, this was not the case for *Acta2*, which was upregulated in non-regenerating P8 and adult mouse, relatively weakly upregulated in regenerating P1 neonatal mouse, and low/absent in zebrafish. Interestingly, in agreement with a very recent study (*54*), we found that the presence Acta2^+^ interstitial cells was significantly increased when regeneration was blocked in zebrafish hearts. Although these cells were not labelled by the *col1a2:mCherry* reporter, Allanki *et al* have revealed that, at least in the cryoinjury model, a proportion of them were of epicardial origin. A relative lack of Acta2 upregulation in zebrafish interstitial cells during regeneration could indicate a reduction in myofibroblast differentiation, or the nature of this transition, and potentially the associated fibrotic response. Another possibility involves the ability of ACTA2 to enable myofibroblasts to contract during the process of wound healing(*55*). Indeed, wound contraction following injury serves to decrease the amount of tissue which needs to be repaired. In the heart this would allow the damaged tissue to be repaired rapidly in order to avoid rupture, a process which most likely supersedes regeneration in adult mammals. It is therefore conceivable that in regenerating tissue, wound contraction may not be required and could even inhibit regeneration by impeding certain regenerative processes, such as neovascularization, which may benefit from an open/relaxed wound environment. Further research will be required in order to determine whether the lack of Acta2 observed in zebrafish fibroblasts can be linked to reduced myofibroblast differentiation and fibrosis, or whether this affects a specific feature of myofibroblasts, such as contractility.

Our data also highlights that the choice of genetic marker used to isolate and study fibroblasts is critical. Indeed, we found that genes that have previously been reported to be upregulated in fibroblasts in zebrafish, such as *rspo1*(*9*) were in fact expressed by *tcf21^-^* neural crest derivatives. The origin of *rspo1^+^;tcf21^-^* cells in the epicardial layer and whether they play a role in cardiac regeneration will require further investigation.

Following cardiac injury in adult zebrafish there is a rapid endothelial response resulting in wound neovascularization. This precedes the expansion of proliferating cardiomyocytes which will repopulate and ultimately regenerate the missing tissue. Our data indicates that *tal1* plays an essential role in the endothelial regenerative response and that Tal1 inhibition, for example by the expression of a dominant negative isoform as we have shown here, impedes cardiac regeneration. Tal1 has been linked to a number of endothelial processes which may affect the regenerative response, such as the regulation of endocardial cell-cell contacts, endocardial identity and neovascularization(*56, 57*). Although *tal1* expression does not increase dramatically during cardiac regeneration, based on previous research it is reasonable to speculate that the genetic programs that Tal1 regulates are, very likely, dictated by its co-factors(*35*). Indeed, a combination of chromatin immunoprecipitation (ChIP) and transcriptomic analysis has shown that, in the absence of *Lmo2*, TAL1 relocates to different DNA target sites where it regulates alternative genetic programs. Our scRNA-seq data indicates that the expression of the Tal1 co-factor, *lmo2*, decreases in *tal1^+^* endothelial cells at 7dpa. Intriguingly, we also observed an increase in expression of the tight junction associated gene *cgnl1* at 7dpa in *tal1^+^* endothelial cells. *Cgnl1* has also been implicated in neovascularization(*58*). Because Tal1 has previously been described to regulate endothelial tight junctions during endocardial development it is tempting to speculate that this process has been disrupted following Tal1 inhibition, resulting in defective cardiac regeneration.

Macrophages are key regulators of regeneration and evidence for their involvement in cardiac regeneration has been demonstrated in a variety of different species(*12, 15, 59*). Our scRNA-seq data has revealed a number of interesting features associated with macrophages during cardiac regeneration. Firstly, although resident macrophages in the uninjured heart appear to have a more pro-inflammatory signature compared to the recruited 3dpa population we observed, we cannot rule out the possibility that the situation may be reversed at an earlier timepoint, as appears to be the case in regenerating neonatal mouse hearts. Previous reports have indicated that during regeneration in neonatal mice there is no recruitment of pro-inflammatory *Ccr2* expressing macrophages(*60*). However, analysis of published scRNA-seq data indicates a rapid influx of *Ccr2* expressing macrophages 1 day post MI coincident with elevated *Tnfa* and *Il1b* expression, an event which may have been missed previously. Furthermore, at 3 days post MI in regenerating neonatal mouse hearts, despite being more numerous, macrophages expressed lower levels of *Tnfa* and *Il1b* compared to sham operated animals. We observed a similar increase in macrophage cell number but reduction in *tnfa* and *il1b* expression within these cells in zebrafish hearts 3 days after injury. These data suggest that the inflammatory response of macrophages in regenerating adult zebrafish hearts is more reminiscent of the regenerating neonatal mouse macrophage response, however future studies at earlier timepoints will be required to confirm these observations. Interestingly, in regenerating zebrafish hearts we could also detect a population of proliferating macrophages. Similarly, proliferating macrophages have been observed in mice post MI which serves to maintain macrophage numbers during the injury response(*42*). More focused studies will be required in order to determine whether the initial macrophage response, occurring during the first hours/days after injury, reflects any difference in regenerative capacity between adult zebrafish and mammals.

Our data also indicates that zebrafish possess a potentially unique mechanism for regulating the macrophage response to CXCL chemokine signals. CXCR3 and its ligands are responsible for recruiting macrophages to sites of injury/infection(*61*). Previous research indicates that zebrafish possess 2 *CXCR3* orthologs, *cxcr3.2*, which is a functional G-protein coupled receptor, and *cxcr3.3* which lacks downstream signaling capabilities. It is apparent that Cxcr3.3 acts as ligand scavenger, reducing the amount of Cxcr3 ligands available to bind and activate Cxcr3.2(*18*). Our data indicates that during cardiac regeneration in zebrafish, all macrophages express *cxcr3.2* in relatively equal proportions, however, resident macrophages also express nearly 2 fold more *cxcr3.3* than recruited macrophages and as such it is fair to assume that this will reduce their ability to respond to Cxcr3 ligands. This may provide zebrafish with an elegant mechanism for fine-tuning the resident M1 macrophage response to Cxcl ligands. For example, modulating Cxcr3 signalling may play a role in blunting the inflammatory response of the M1 resident macrophages during cardiac regeneration. It could also potentially serve to reduce the mobility of resident macrophages and help to maintain this population within the heart.

MMPs have frequently been associated with cardiac injury and regeneration. In particular, MMP14 appears to be the major type 1 collagenase in ischemic mouse hearts(*62*). Our data indicates that Mmp14b is particularly enriched in recruited macrophages at 3dpa. Furthermore, we observed that inhibiting the collagenolytic activity of Mmp14 resulted in defective migration of macrophages into the injury site and a subsequent failure to regenerate the myocardium, leading to the formation of large collagen/fibrin scar. Our scRNA-seq analysis of Mmp14 inhibited macrophages indicated that even at this early time point the expression of a number of genes associated with the regenerative response have become misregulated. Of potential interest is HB EGF which has been linked to a variety of processes important for cardiac regeneration(*63, 64*). However, expression of *hbegfb* appears to increase when cardiac regeneration fails following Mmp14 inhibition. In this context HB EGF has previously been shown to have detrimental effects on the remodeling process after myocardial infarction in mammals by enhancing fibroblast activation and invasion(*65*). It could therefore play a role in the development of the substantial fibrin/collagen scar that is present at 30dpa when Mmp14 is inhibited. Re-analysis of scRNA-seq data from adult mice and neonates following myocardial infarction showed that, conversely to what we observed in zebrafish, *Mmp14* expressing were not present in the myocardium prior to injury. Whether the presence of Mmp14b-producing resident macrophages confers zebrafish with some kind of regenerative advantage, perhaps by allowing resident macrophages to respond in a similar manner to recruited macrophages, will require further investigation. However, in stark contrast to our own observations in zebrafish, it appears that in adult mice, MMP14 plays a deleterious role after cardiac injury(*62*). Indeed MMP14 heterozygote KO mice display a marked improvement in survival post MI due to reductions in infarct size, left ventricular dilation and compensatory hypertrophy. Furthermore, there is also a significant reduction in the number of macrophages localised to the infarct area in MMP14 +/- mice, similar to our own observations in regenerating zebrafish hearts when Mmp14 is inhibited. More recently, it has been shown that macrophage specific deletion of MMP14 in adult mice reduces left ventricular dysfunction following MI(*66*). It appears that loss of MMP14 in macrophages results in attenuated, Tgfβ dependent, fibrosis. Our own data indicated that *mmp14b* expressing macrophages are recruited to the wound site a few days after injury, similar to observations in mouse MI models. Targeting Mmp14 in zebrafish with a specific inhibitor disrupted cardiac regeneration, which appears to be the complete inverse of the situation described in adult mice. Why this is the case remains unclear. It is possible that zebrafish Mmp14b possesses properties that are absent from mammalian MMP14, or this could be due to the expression of *mmp14b* by resident macrophages in uninjured zebrafish hearts, a feature that is not present in neonatal or adult mouse hearts. It may also be the case that MMP14 macrophages are performing a similar role in adult mouse hearts following injury but that another pro-regenerative process is absent. A failure to co-ordinate a multi-faceted regenerative response could in fact be detrimental. Future research will be required to determine why MMP14 plays a positive pro-regenerative role in adult zebrafish, and yet appears to exacerbate the damage associated with MI in mammals.

In summary our data has highlighted a number of intriguing features of the interstitial cellular response during cardiac regeneration in adult zebrafish. Although there are notable differences when compared to non-regenerating mammalian hearts there are also many similarities which certainly offers hope that we will eventually be able to recapitulate this process in adult humans.

## Materials and methods

### Single cell RNA sequencing

Hearts were dissected, placed in cold HBSS with heparin. Atria and outflow tracts removed, ventricles opened and washed. 10 ventricles were pooled for each sample: unamputated, sham-operated, 3 dpa, 3 dpa +vehicle, 3 dpa+inhibitor, 7 dpa and 14 dpa. Cells were pre-incubated for 2h on ice in HBSS-7% TrypLE. HBSS TrypLE was removed and tissue was dissociated in HBSS Collagenase (type II, 5mg/ml; type IV, 5mg/ml) with CaCl2 (12μM) for 1h on a rotator (800RPM) at 32 degree. Dissociated hearts were put through a 40μm cell strainer, spun down (300g, 5min at 4 degrees) and recovered in FACS buffer (HBSS 2% FCS). Nuclei were stained by adding Vybrant DyeCycle Ruby Stain to the cell suspension and heated for 10min at 32 degrees before adding DAPI and sorting into PBS 0.04% BSA. Sorting was performed on a BD FACS Melody.

**Supplemental Table I.**

| Sample | unamputated<br>control | 3 days post-amputation | 7 days post-amputation | 14 days post-amputation |
| --- | --- | --- | --- | --- |
| number of hearts<br>pooled/dissociated | 10 | 10 | 10 | 10 |
| number of cells | 1 876 | 1 565 | 3 153 | 2 475 cells |
| median genes per<br>cell | 1 411 | 1 339 | 1 425 | 1 450 |

Cellular suspensions were loaded on a Chromium controller (10x Genomics, Pleasanton, CA, USA) to generate single-cell Gel Beads-in-Emulsion (GEMs). Single-cell RNA-Seq libraries were prepared using Chromium Single cell 3’RNA Gel Bead and Library Kit v3.1. GEM-RT was performed in a C1000 Touch Thermal cycler with 96-Deep Well Reaction Module (Bio-Rad): 53°C for 45 min, 85°C for 5 min; held at 4°C. After RT, GEMs were broken and the single-strand cDNA was cleaned up with DynaBeads MyOne Silane Beads (Thermo Fisher Scientific). cDNA was amplified using the C1000 Touch Thermal cycler with 96-DeepWell Reaction Module: 98°C for 3 min; cycled 12: 98°C for 15 s, 63°C for 20 s, and 72°C for 1 min; 72°C for 1 min; held at 4°C. Amplified cDNA product was cleaned up with the SPRI select beads. Indexed sequencing libraries were constructed following these steps: (1) fragmentation, end-repair and A-tailing and size selection with SPRIselect; (2) adapter ligation and cleanup with SPRIselect; (3) sample index PCR and size selection with SPRIselect. The barcoded sequencing libraries were quantified by quantitative PCR (KAPA Biosystems Library Quantification Kit for Illumina platforms). Sequencing libraries were loaded at 300 pM on an Illumina NovaSeq6000 using the following read length: 28 bp Read1, 8 bp I7 Index, 91 bp Read2 (experiment 1), and 28 bp Read1, 10 bp I7 Index, 10 bp I5 Index, 87 bp Read2 (experiment 2).

Image analyses and base calling were performed using the NovaSeq Control Software and the Real-Time Analysis component (Illumina). Demultiplexing was performed using the 10X Genomics software Cellranger mkfastq (v3.1.0 for experiment 1 and v6.0.1 for experiment 2), a wrapper of Illumina’s bcl2fastq (v2.20). The quality of the raw data was assessed using FastQC (v0.11.8) from the Babraham Institute and the Illumina software SAV (Sequencing Analysis Viewer). FastqScreen (v0.14.0) was used to estimate the potential level of contamination.

Alignment, gene expression quantification and statistical analysis were performed using Cell Ranger count on *Danio rerio* ’s transcriptome GRCz11 (sequences and annotation were downloaded from Ensembl! on July 24th, 2019). In order to discard ambient RNA falsely identified as cells, Cell Ranger count was run a second time with the option --force-cells to force the number of cells to detect. Cell Ranger aggr was then used to combine each sample result into one single analysis.

### Data availability

Datasets generated for this study: sequencing data have been deposited in the ArrayExpress database at EMBL-EBI (www.ebi.ac.uk/arrayexpress) under accession code E-MTAB-10643.

Previously published datasets used for this study: E-MTAB-7376 (www.ebi.ac.uk/arrayexpress) from Farbehi et al.(*6*); GSE153480 (www.ncbi.nlm.nih.gov/geo/) from Wang et al.(*7*).

### Zebrafish transgenic lines and husbandry

Zebrafish were maintained under standardized conditions and experiments were conducted in accordance with local approval (APAFIS#4054-2016021116464098 v5) and the European Communities council directive 2010/63/EU. Embryos were staged as described (*67*). The *Tg(fli1a:GFP)y1Tg* was provided by the CMR[B] *Centro de Medicina Regenerativa de Barcelona. Tg(gata1:dsred)* and *Tg(mpeg1.1:mCherry)* were provided by the Lutfalla Lab, University of Montpellier. *Tg(col1a2:mCherry)* was provided by the Mercader Lab, Bern University. *Tg(eab2:[EGFP-T-mCherry])^vu295^* was provided by the Chen Lab, Vanderbilt University Medical Center. All larvae and adults were euthanised by administration of excess anaesthetic (Tricaine).

### Zebrafish cardiac regeneration

All amputations were performed as described(*2*), in accordance with local approval (APAFIS#4052). For the scRNA-seq analysis we used 6 month old sibling offspring from an incross of *Tg(cmlc2a:GFP)* which were generated on an AB wildtype background.

### Neonatal heart regeneration

All amputations were performed on Swiss-JL (Janvier labs) P1 neonatal mice as described(*68*),in accordance with local approval (APAFIS#1498-15516).

### Edu labelling

EdU labelling was performed according to the manufacturers instructions (Click-iT EdU Kit C10337, Molecular Probes). Amputated adult *Tg(mpeg1.1:mCherry^+^)* were anesthetized in Tricaine then injected with 50μl of a 240μg/ml EdU solution daily.

### Immunohistochemistry

Immunohistochemistry was performed on 10 μm cryo-sections as previously described(*59*). The primary antibodies used in this manuscript are anti-RFP (5F8 Chromotek), anti-GFP (GFP1020 Aves), anti-Acta2 GTX124505 Genetex), anti-RSPO1 (ab106556 Abcam), anti-Tal11 (abx339062 Abbexa), anti-α Sarcomeric actin (A2172 Sigma), anti-MMP14 (mbs422986 Mybiosource and GTX128198 Genetex), IB4 (Isolectin GS1B4) (I21413 Invitrogen), EdU labelling was performed according to the manufacturers instructions (Click-iT EdU Kit C10337, Molecular Probes).

### Histology

Acid Fuchsin-Orange G (AFOG) staining was performed on 10 μm cryosections as previously described (*69*).

### Imaging

An Olympus SZX16 fluorescence stereomicroscope fitted with a Jenoptik ProgRes CF Cool CCD Microscope Camera was used for histological section imaging and a Leica TCS SP-8 confocal microscope for imaging immunohistochemical labelling. Image acquisition and image analysis were performed on workstations of the Montpellier RIO Imaging facility of Arnaud de Villeneuve.

### Dominant negative Tal transgenic zebrafish

The DN *tal* construct and transgenic line were generated using the Tol2 Kit as described (*70, 71*). Dominant negative zf *tal1* was generated as described(*36*). For the *Tg(fliEP:loxRFPlox:DNtal*) construct the 5’ entry clone 478 p5Efli1ep was a gift from Nathan Lawson (Addgene plasmid # 31160; http://n2t.net/addgene:31160; RRID:Addgene_31160, the middle entry clone contained a floxed RFP stop cassette amplified from pBOB-LRL-CBReGFPpA (a kind gift from Geoff Whal) and the 3’ entry clone contained zebrafish dominant negative *tal1*. For the *Tg*(*fliEP:Ert2CreErt2)* the 5’ entry clone was p5Efli1ep and the middle entry clone was pMEErt2CreErt2 as described(*2*).

### Real-time quantitative RT-PCR

RNA was extracted from amputated/unamputated ventricles of AB wild type zebrafish and quantitative PCR was performed using a Roche LightCycler 480 system as described (*34*). Primer sequences are provided below.

MMP2 forward- 5’ GGTGTGCAACCACTGAAGAT 3’

MMP2 reverse- 5’ AGGGTGCTCCATCTGAATTT 3’

MMP9 forward- 5’ TTTGACGCCATCACTGAAAT 3’

MMP9 reverse- 5’ TTCGCAGAGATCATGAAAGG 3’

MMP13a forward- 5’ CTCAGAGCCCAGATGTTGAA 3’

MMP13a reverse- 5’ CCTTCTCACCTTTGATCAGGA 3’

MMP14a forward- 5’ CTCGCAAGTGTGTTTCTGGT 3’

MMP14a reverse- 5’ TCACCAGGAGGAAGATACCC 3’

MMP14b forward- 5’ GATATGAAACCTGAGGCATGG 3’

MMP14b reverse- 5’ GTGACTGTCAGGCCGTAGAA 3’

### Chemical treatments

Transgenes were expressed by inducing Cre mediated recombination with tamoxifen as described(*2*). GM6001 (ab120845, Abcam) was dissolved in DMSO to reach a stock concentration of 10 mM and then split into 10 μl aliquots and stored at – 20°C. On the day of injection, the 10 μl aliquot was added to 1 ml 1X PBS to reach a final concentration of 100 μM. This solution was administered daily i.p. Control groups were administered daily with 1X PBS. NSC40520 (SML0518 Sigma) was diluted in DMSO to reach a stock concentration of 10 Mm. 5 fish were placed in a beaker with 500ml of system water and 500μl of NSC40520 was added to reach a final concentration of 10 μM. Controls were placed in a beaker and 500μl of DMSO was added. The fish were left overnight then rinsed the next morning and returned back to the main aquarium during the day. This was repeated for the duration of each experiment.All chemical treatments were performed in accordance with local approval (APAFIS#4054).

### Statistical analysis

GraphPad prism was used to perform all statistical analysis. Details of statistical analysis are provided in figures and figure legends.

## Acknowledgements.

We acknowledge the imaging facility MRI, member of the national infrastructure France-BioImaging infrastructure supported by the French National Research Agency (ANR-10-INBS-04, «Investments for the future»). Montpellier Genomix (MGX). MGX acknowledges financial support from France Génomique National infrastrusture, funded as part of “Investissement d’Avenir” program managed by Agence National pour la Recherche (contract ANR-10-INBS-09). The Jopling lab is part of the Laboratory of Excellence Ion Channel Science and Therapeutics supported by a grant from the ANR. Work in the Jopling lab is supported by a grant from the “la Fondation Leducq” and from the ANR (contract ANR-20-CE14-003 MetabOx-Heart).

---

**Supplemental Figure 1.**
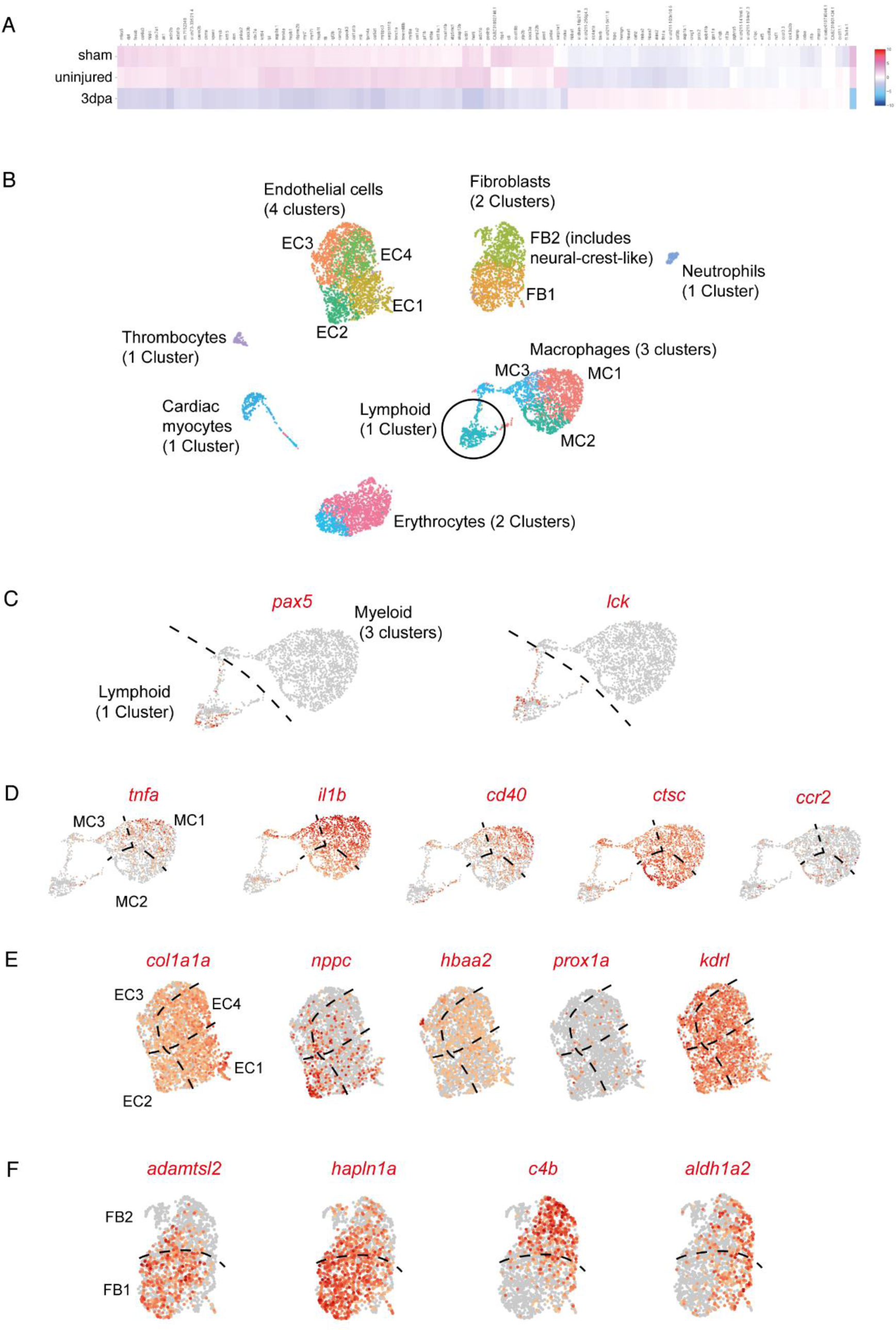
Single cell RNAseq analysis of regenerating zebrafish hearts. (**A**) Heat map depicting differentially expressed genes among sham operated, uninjured and 3dpa samples. (**B**) UMAP plot highlighting the different clusters within in each population of cells identified in regenerating zebrafish hearts. In particular, 4 clusters were identified within the endothelial population of cells (EC1-4), 2 clusters were identified within the epicardium/fibroblast population of cells (FB1,2) and 3 clusters were identified within the myeloid/macrophage population of cells (MC1-3). (**C-F**) UMAP plots depicting the relative expression of particular genes within the different sub clusters. (**C**) Lymphoid cells (*pax5* and *lck*). (**D**) Macrophages (*tnfa*, *il1b*, *cd40*, *ctsc*, *ccr2*). (**E**) Endothelium (*col1a1a*, *nppc*, *hbaa2*, *prox1a*, *kdrl*). (**F**) Fibroblasts (*adamtsl1*, *hapln1a*, *c4b*, *aldh1a2*).

**Supplemental Figure 2.**
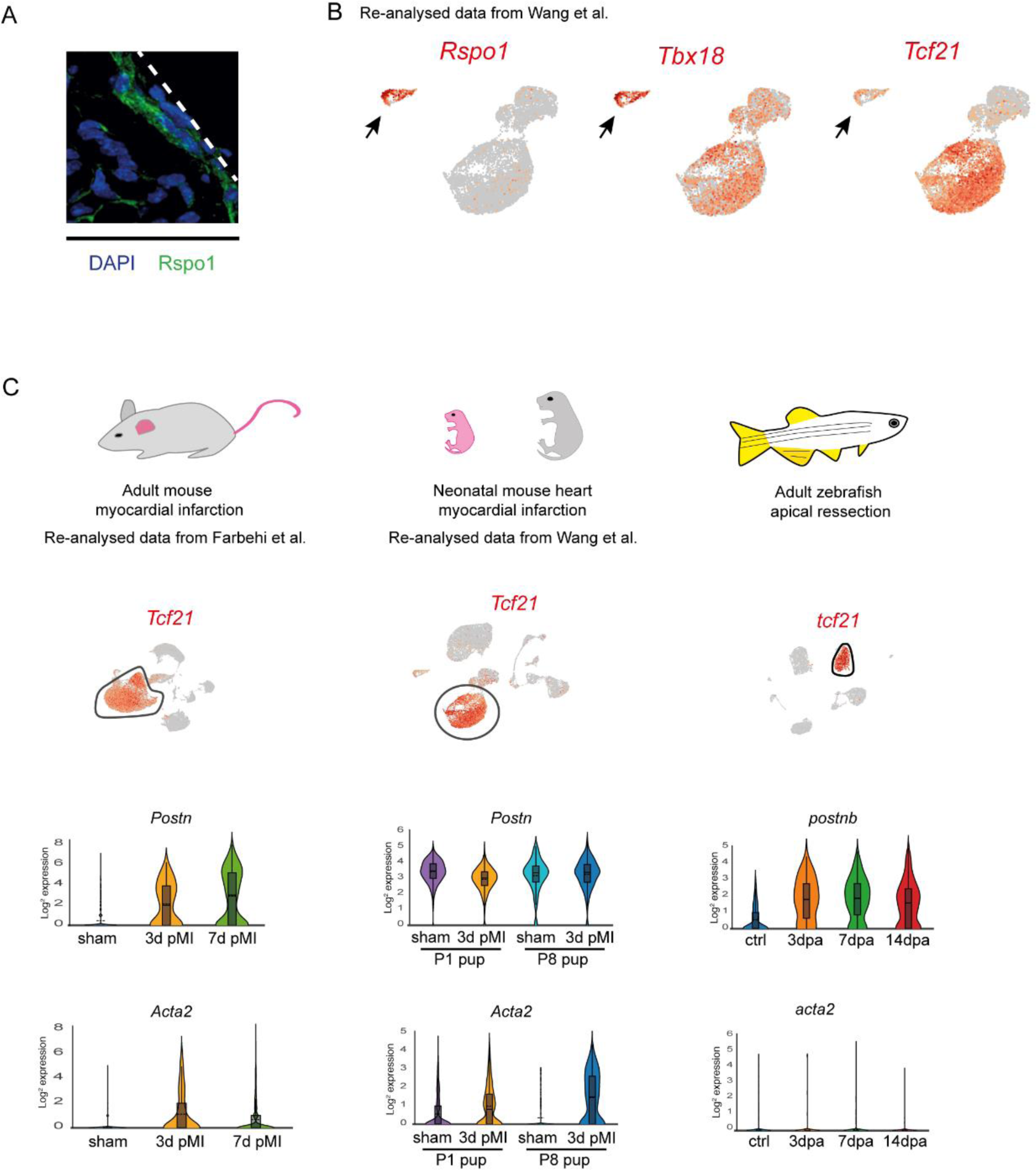
Mesenchymal lineages and cardiac fibroblast activation in the regenerating zebrafish heart. (**A**) IHC image of 7dpa neonatal mouse heart labelled with RSPO1 antibody (green), white dashed line indicates the outer edge of the epicardium. (**B**) UMAP plots from re-analysed neonatal mouse scRNA-seq data (*Wang et al.)* depicting the co-expression of *Rspo1* with *Tbx18* and *Tcf21*. Note that although *Rspo1* is co-expressed with both genes in a smaller cluster (black arrow) there is relatively little expression within the main *Tcf21^+^* fibroblast cluster. (C) Re-analysis of scRNA-seq data from adult (Farbehi *et al.)* and neonatal mice (Wang et al.) after myocardial infarction compared with adult zebrafish after apical resection. Adult mouse data is from 3 and 7 days post myocardial infarction (MI). Neonatal is from postnatal day 1 and postnatal day 8 pups, with sham and 3 days post MI samples. UMAP plots highlight the fibroblast population (*Tcf21*) in each dataset. Violin plots represent the expression of *Periostin* and *Acta2* at different time points after injury in each dataset. Note-*Periostin* is expressed in uninjured neonatal mouse hearts, *Acta2* is expressed by fibroblasts in both adult and neonatal mice after injury but not by adult zebrafish fibroblasts.

**Supplemental Figure 3.**
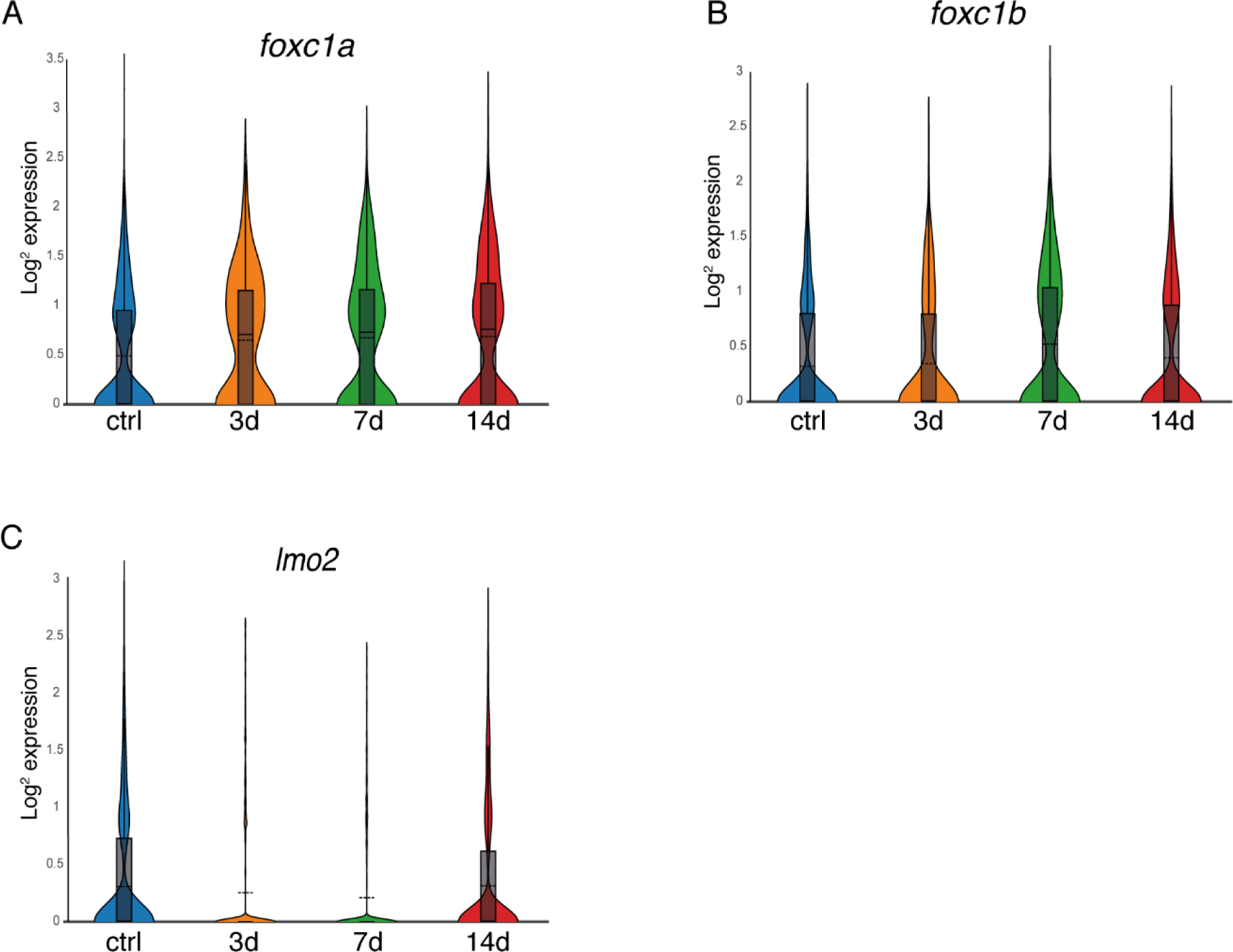
Tal1 is a regulator of the endothelial regenerative response. (**A-C**) Violin plots comparing the expression of *foxc1a* (**A**), *foxc1b* (**B**) and *lmo2* (**C**) at different time points after injury (uninjured (ctrl), 3dpa, 7dpa and 14dpa) within the endothelial population of cells.

**Supplemental Figure 4.**
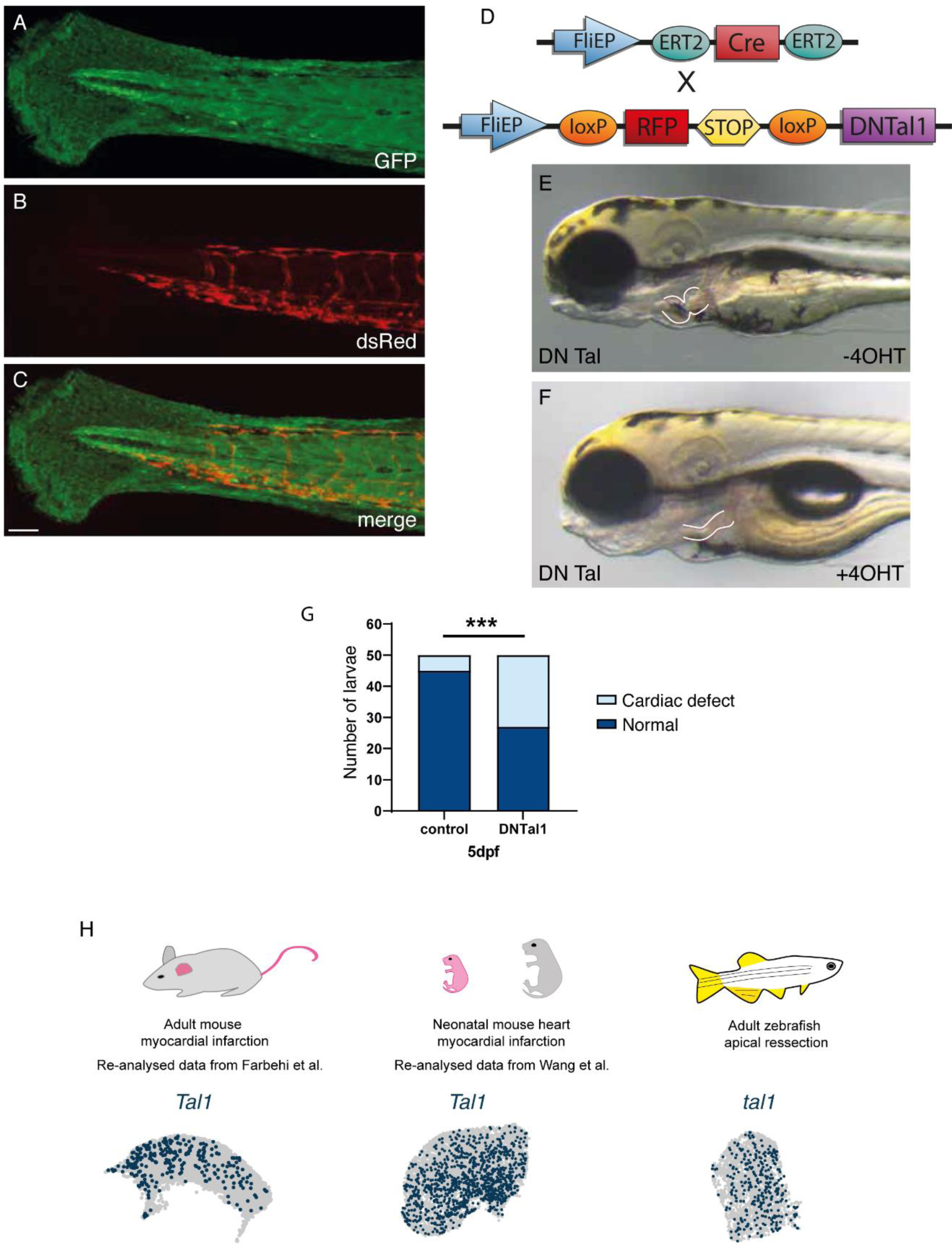
Tal1 is a regulator of the endothelial regenerative response. (**A-C**) IHC images of a *Tg(eab2:[EGFP-T-mCherry])vu295* / *Tg*(*fliEP:Ert2CreErt2)* 5dpf larvae treated with tamoxifen. (**A**) GFP is ubiquitously expressed. (**B**) DsRed is expressed in endothelial cells following Cre mediated recombination. (**C**) Merged image of A and B. Note the vasculature is labelled with dsRed (**B**) indicating that Ert2CreErt2 in *Tg*(*fliEP:Ert2CreErt2)* zebrafish is restricted to endothelial cells. (**D**) Diagram indicating the 2 constructs used to generate the *Tg*(*fliEP:Ert2CreErt2;fliEP:loxRFPlox:DNtal*) transgenic zebrafish line. (**E,F**) *Tg*(*fliEP:Ert2CreErt2;fliEP:loxRFPlox:DNtal*) transgenic zebrafish larvae either untreated -4OHT (**E**) or tamoxifen treated +4OHT (**F**). The developing heart has been highlighted with white lines, note that the heart in the tamoxifen treated larvae has failed to develop normally. (**G**) Graph depicting the number of larvae which develop a cardiac defect in either untreated (control) or tamoxifen treated (DN Tal1) groups. Fisher’s exact test, n=50 larvae for each condition (***: p<0.001). (**H**) Re-analysis of scRNA-seq data from post myocardial infarction adult mice (Farbehi *et al*.) and neonatal mice (Wang *et al*.) compared with adult zebrafish after apical resection. UMAP plots indicating the expression of *Tal1* (dark blue dots) within the endothelial population of cells in each dataset.

**Supplemental Figure 5.**
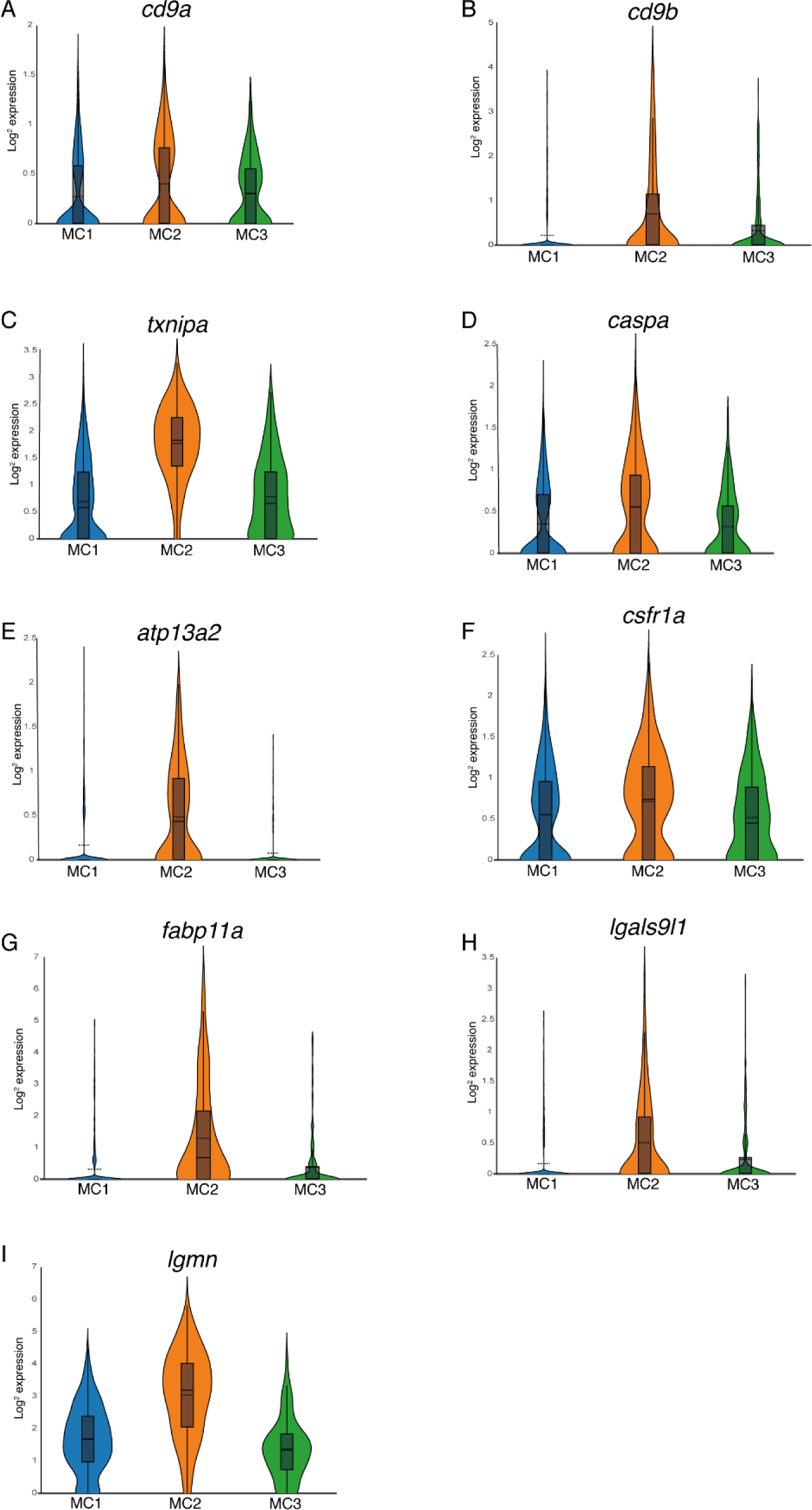
Genes enriched in the recruited macrophage population. (**A-I**) Violin plots comparing the expression of different genes within the macrophage subpopulations (MC1-resident macrophages, MC2-recruited macrophages and MC3-proliferating macrophages), *cd9a* (**A**), *cd9b* (**B**), *txnipa* (**C**), *caspa* (**D**), *atp13a2* (**E**), *csfr1a* (**F**), *fabp11a* (**G**), *lgals9l1* (**H**) and *lgmn* (**I**).

**Supplemental Figure 6.**
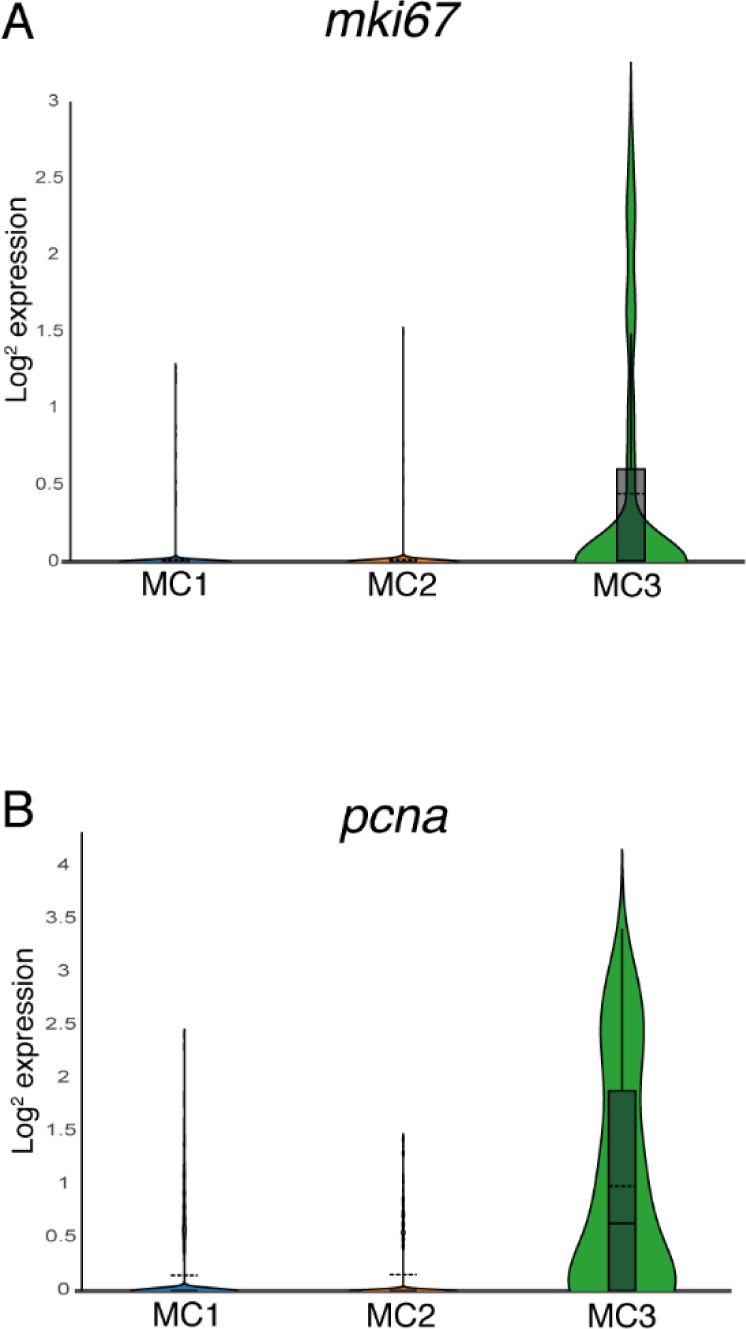
Genes enriched in the proliferating macrophage population. (**A,B**) Violin plots comparing the expression of different genes within the macrophage subpopulations (MC1-proliferating macrophages, MC2-recruited macrophages and MC3-resident macrophages), *mki67* (**A**) and *pcna* (**B**).

**Supplemental Figure 7.**
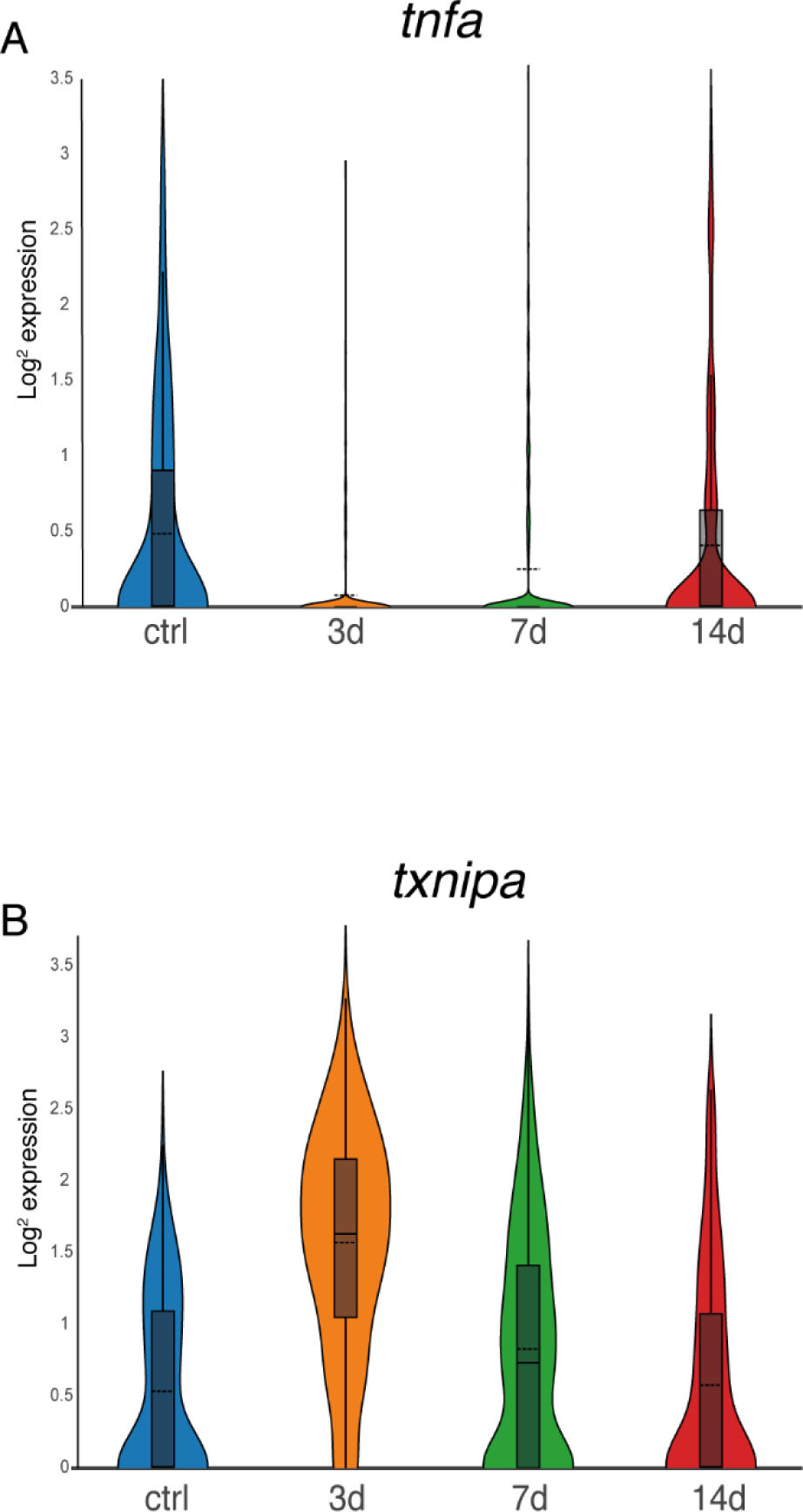
Dynamic expression of inflammatory genes in macrophages at different time points during cardiac regeneration. (**A,B**) Violin plots comparing the expression of different genes by macrophages at different time points during cardiac regeneration (uninjured (ctrl), 3dpa, 7dpa and 14dpa), *tnfa* (**A**), and *txnipa* (**B**).

**Supplemental Figure 8.**
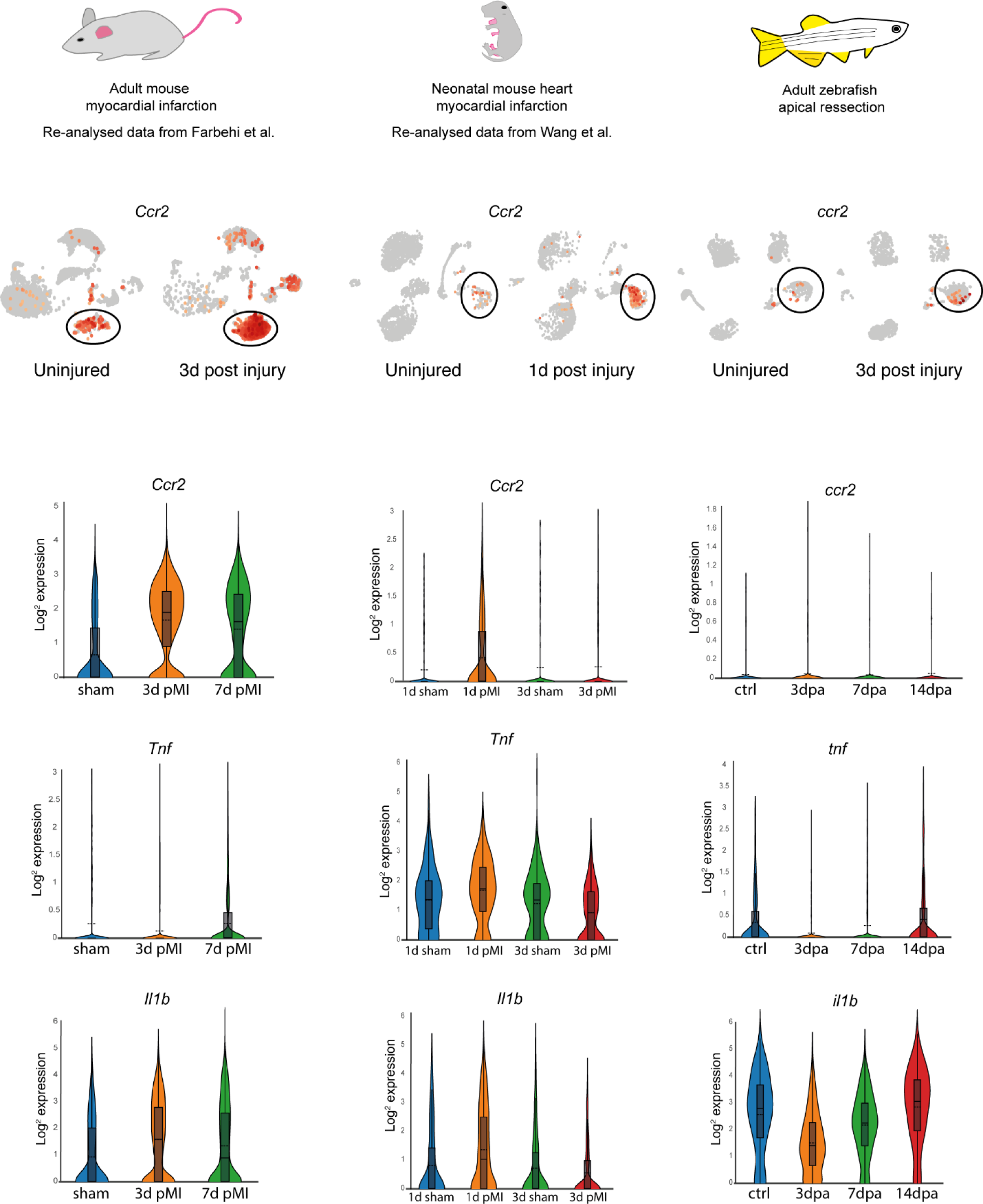
Comparison of recruited macrophages between species. *Adult mouse myocardial infarction.* Re-analysed UMAP plots depicting the expression of *Ccr2* in uninjured sham conditions vs 3 days post injury. Black circles indicate the macrophage population. Violin plots showing the expression of *Ccr2*, *Tnfa* and *Il1b* at different time points after injury (sham, 3 days post myocardial infarction (d pMI) and 7d pMI) in macrophages. *Neonatal mouse myocardial infarction.* Re-analysed UMAP plots depicting the expression of *Ccr2* in uninjured sham conditions vs 1 day post injury. Black circles indicate the macrophage population. Violin plots showing the expression of *Ccr2*, *Tnfa* and *Il1b* at different time points after injury (1 day sham, 1 day post myocardial infarction (d pMI) and 3d sham, 3d pMI) in macrophages. *Adult zebrafish apical resection*. UMAP plots depicting the expression of *Ccr2* in uninjured control conditions vs 3 days post injury. Black circles indicate the macrophage population. Violin plots showing the expression of *Ccr2*, *Tnfa* and *Il1b* at different time points after injury (control, 3 days post amputation (dpa) and 7dpa and 14dpa) in macrophages.

**Supplemental Figure 9.**
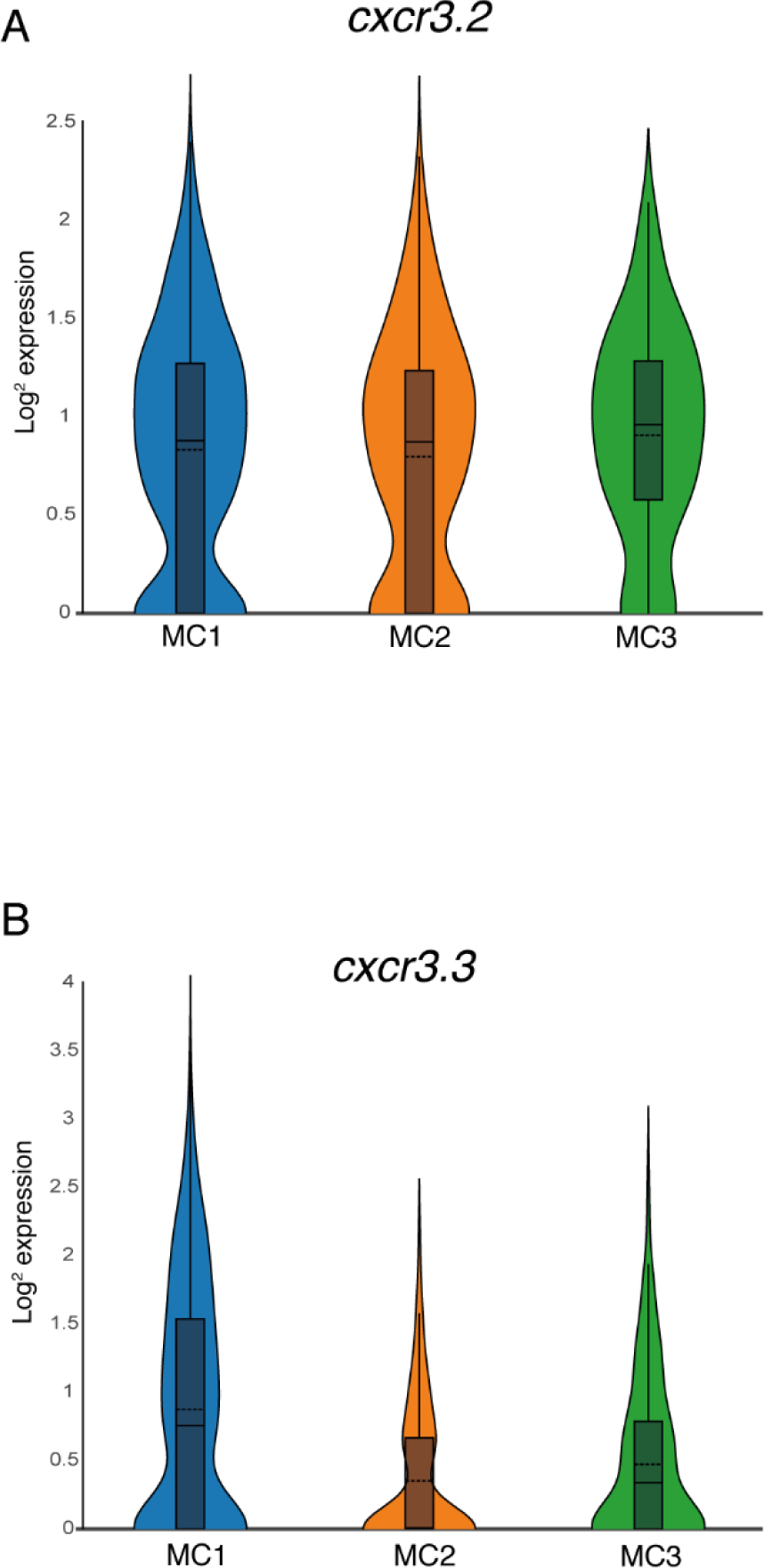
The expression of *cxcr3.2* and *cxcr3.3* within the macrophage subpopulations. (**A,B**) Violin plots comparing the expression of *cxcr3.2* and *cxcr3.3* within the macrophage subpopulations (MC1-resident macrophages, MC2-recruited macrophages and MC3-proliferating macrophages), *cxcr3.2* (**A**) and *cxcr3.3* (**B**).

**Supplemental Figure 10.**
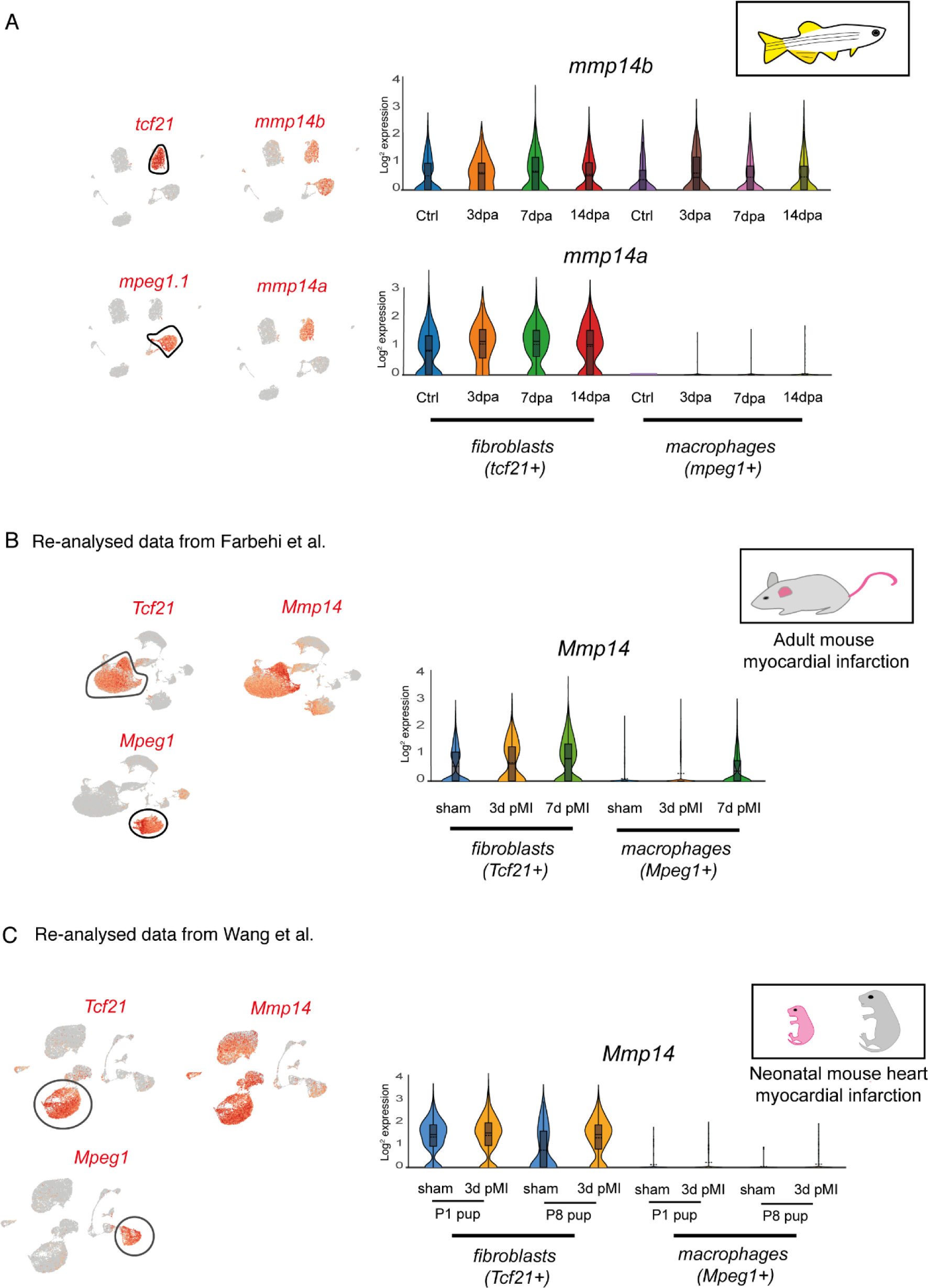
Comparison of Mmp14 expression in macrophages between species. (**A**) Adult zebrafish apical resection. UMAP plots depicting the expression of *mmp14b* and *mmp14a* in relation to *tcf21* expressing fibroblasts and *mpeg1.1* expressing macrophages. Note *mmp14b* is expressed by fibroblasts and macrophages while *mmp14a* is restricted to fibroblasts. Violin plots showing the expression of *mmp14b* and *mmp14a* at different time points during regeneration (uninjured (ctrl), 3dpa, 7dpa and 14dpa) in fibroblasts and macrophages. (**B**) Adult mouse myocardial infarction. Re-analysed UMAP plots depicting the expression of *Mmp14* in relation to *Tcf21* expressing fibroblasts and *Mpeg1* expressing macrophages. Violin plots showing the expression of *Mmp14* at different time points after injury (sham, 3 days post myocardial infarction (d pMI) and 7d pMI) in fibroblasts and macrophages. Note that *Mmp14* expression appears at 7d pMI in the macrophage population. (**C**) Neonatal mouse myocardial infarction. Re-analysed UMAP plots depicting the expression of *Mmp14* in relation to *Tcf21* expressing fibroblasts and *Mpeg1* expressing macrophages. Violin plots showing the expression of *Mmp14* at different time points after injury (sham, 3d pMI) by fibroblasts and macrophages in regenerating P1 pups and non-regenerating P8 pups. Note that *Mmp14* expression is absent in the macrophage population.

**Supplemental Figure 11.**
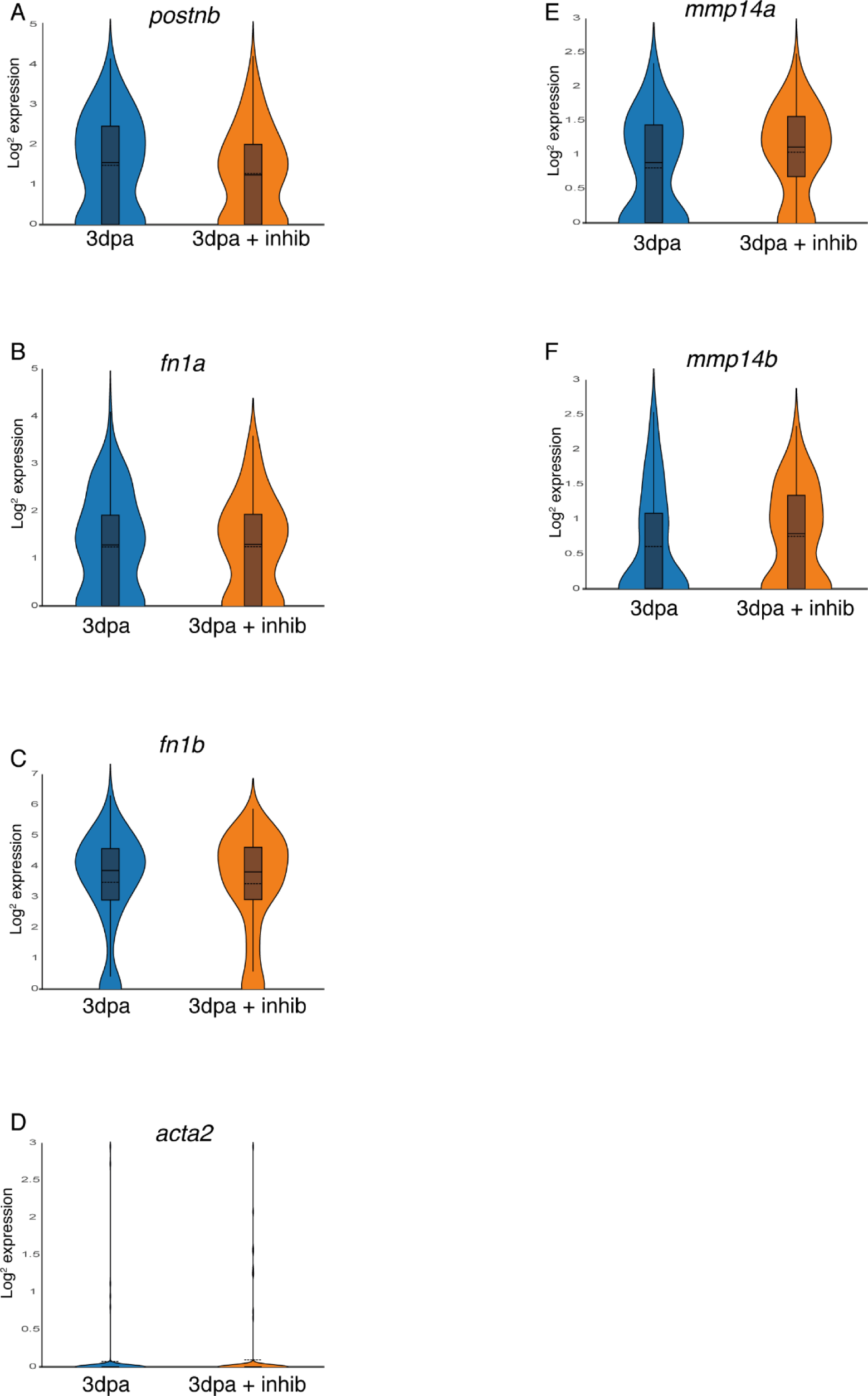
Comparison of gene expression by fibroblasts following Mmp14 inhibition. (**A-F**) Violin plots comparing the expression of different genes within the fibroblast population in untreated (ctrl) or MMP14 inhibitor treated (+inhib) 3dpa regenerating zebrafish hearts, *postnb* (**A**), *fn1a* (**B**), *fn1b* (**C**), *acta2* (**D**), *mmp14a* (**E**) and *mmp14b* (**F**).

**Supplemental Figure 12.**
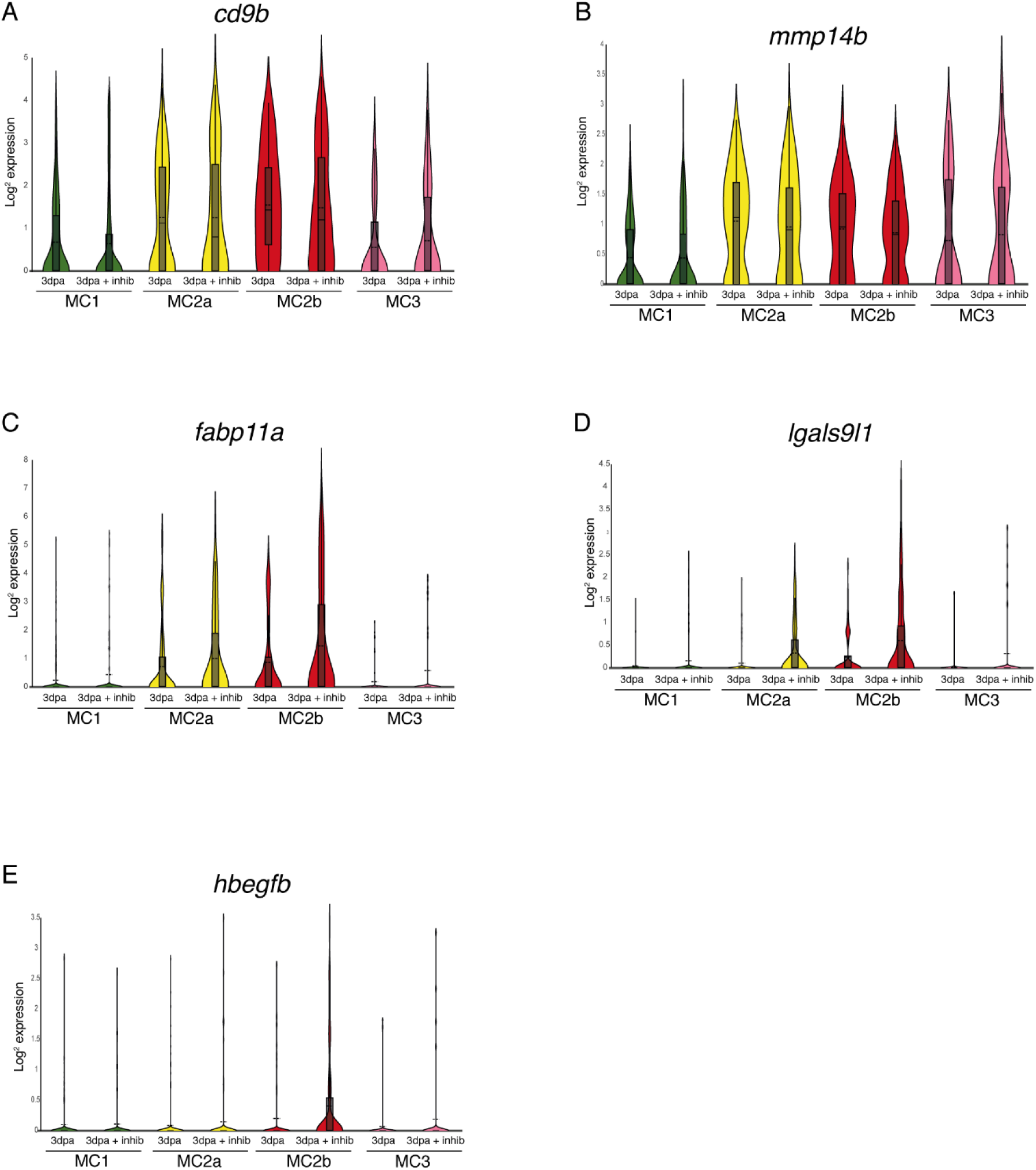
Comparison of gene expression within the macrophage subpopulations following Mmp14 inhibition. (**A-E**) Violin plots comparing the expression of different genes within the macrophage subpopulations (MC1, MC2a, MC2b and MC3) in untreated (ctrl) or MMP14 inhibitor treated (+inhib) 3dpa regenerating zebrafish hearts, *cd9b* (**A**), *mmp14b* (**B**), *fabp11a* (**C**), *lgals9l1* (**D**), and *hbegfb* (**E**).

